# Phage-induced efflux down-regulation boosts antibiotic efficacy

**DOI:** 10.1101/2023.09.21.558807

**Authors:** Samuel Kraus, Urszula Łapińska, Krina Chawla, Evan Baker, Erin L Attrill, Paul O’Neill, Audrey Farbos, Aaron Jeffries, Edouard E. Galyov, Sunee Korbsrisate, Kay B. Barnes, Sarah V Harding, Krasimira Tsaneva-Atanasova, Mark A T Blaskovich, Stefano Pagliara

## Abstract

The interactions between a virus and its host vary in space and time and are affected by the presence of molecules that alter the physiology of either the host or the virus. Determining the molecular mechanisms underpinning these interactions is paramount for predicting the fate of bacterial and phage populations and for designing rational phage-antibiotic therapies. We study the interactions between stationary phase *Burkholderia thailandensis* and the phage ΦBp-AMP1. Although heterogeneous genetic resistance to phage rapidly emerges in *B. thailandensis*, the presence of phage enhances the efficacy of three major antibiotic classes, the quinolones, the beta-lactams and the tetracyclines, but antagonizes tetrahydrofolate synthesis inhibitors. We discovered that enhanced antibiotic efficacy is underpinned by reduced antibiotic efflux in the presence of phage. This new phage-antibiotic therapy allows for eradication of stationary phase bacteria, whilst requiring reduced antibiotic concentrations, which is crucial for treating infections in sites where it is difficult to achieve high antibiotic concentrations.

## Introduction

Antimicrobial resistance (AMR) has a dramatic impact on global health with an estimated 5 million deaths associated with AMR in 2019 alone ^1^. *Burkholderia* Bptm species cause life-threatening diseases ^2–5^ and are challenging to treat with currently available antibiotics as they are intrinsically resistant to aminoglycosides, macrolides and oxazolidinones due to constitutively expressed efflux pumps ^6–9^, whereas an atypical lipopolysaccharide structure plays a crucial role in resistance to cationic peptides such as polymyxins ^2^.

Moreover, misuse and overuse of antibiotics in the food industry, animal husbandry, and medicine, as well as changes in the global environment have recently contributed to the spread of acquired genetic resistance ^10^. *Burkholderia* Bptm species can acquire antibiotic resistance *in vivo* during treatment, which can be fatal if treatment is not shifted to alternative drugs in due course ^2^.

In order to overcome the development of resistance to antibiotic monotherapies, the deployment of two or more different antibiotics in combination therapies has shown some success in treating and preventing infections ^11,12^; however, antibiotics in conventional regimens can antagonize each other ^13,14^. Therefore, several alternatives to traditional antibiotics have been explored in order to treat resistant infections, including: antibody therapy, antimicrobial peptides, probiotics, metal chelation, CRISPR-Cas9, bioengineered toxins, bacteriocins, vaccines and antibodies ^15–19^. Although promising, all of these approaches are limited by the fact that these antimicrobial agents cannot change or adapt in real time.

In contrast, like bacteria, bacteriophages (i.e., viruses that infect bacteria) amplify at the site of an infection and evolve; therefore, also due to their high number and genetic variability, phage constitute a large and valuable reservoir of natural antimicrobials ^20^. However, the use of phage as a monotherapy to treat bacterial infections presents several challenges, particularly narrow host range and the rapid evolution of resistance to phage ^21^. As antibiotics are the current standard of care, using phage as an adjuvant to antibiotics instead of a monotherapy may be a more rational therapeutic use of phage ^22^. Therefore, phage-antibiotic therapy has undergone a robust revitalization in the last seven years ^23–25^.

Enchanced efficacy of combination therapies has historically been attributed to an increase in phage produced from bacteria in the presence of β-lactam antibiotics, relative to production in their absence ^26,27^. More recently, a variety of lytic phage was observed to form larger plaques in the presence of sublethal concentrations of β-lactam, quinolone and tetracycline antibiotics ^28–31^. This effect has been termed phage-antibiotic synergy and has been linked to antibiotic-induced bacterial filamentation which accelerates phage assembly and cell lysis ^28^. However, other recent evidence suggests that phage-antibiotic synergy can be obtained independently of cell filamentation, enhancement of phage production or the strict use of lytic phage, and that temperate phage are also viable for phage-antibiotic therapy if prophages are induced by DNA damaging antibiotics ^32^.

Moreover, phage-antibiotic synergy has mostly been studied with only one or two concentrations of the antimicrobials, which is insufficient to predict combinatorial concentrations that are effective during treatment, leading to mixed results during phage-antibiotic therapy investigations ^22,33^. Indeed, a recent study employing *Escherichia coli* and the lytic phage ϕHP3 tested several orders of magnitude of both phage and antibiotic concentrations, finding that the nature of the interaction between antibiotics and phage depended both on the type and concentration of the antibiotic and phage employed ^34^.

Therefore, it is imperative to discover and understand the molecular mechanisms underpinning the interactions between phage and antibiotics, in order to rationally design successful new phage-antibiotic therapy ^20^. Here we test molecules representative of eight major antibiotic classes in combination with a recently discovered phage, termed ΦBp-AMP1, that infects and kills *Burkholderia pseudomallei* and *Burkholderia thailandensis* ^35–37^. ΦBp-AMP1 is a podovirus with a 45 nm icosahedral capsid, a 20 nm non-contractile tail and a 45 kb genome. In contrast to other *Burkholderia* phage that are strictly lytic ^38^, ΦBp-AMP1 displays a temperature-dependent switch from the temperate (at 25 °C) to the lytic cycle (at 37 °C) ^35–37^. Using optically based microtiter plate assays and genomics, we studied the development of genetic resistance to ΦBp-AMP1 in stationary phase *B. thailandensis*. We investigated the dynamics of the interactions between ΦBp-AMP1 and representative molecules of the major antibiotic classes over multiple orders of magnitude of antibiotic concentrations and phage titers. We determined the molecular mechanisms underpinning these phage-antibiotic interactions via single-cell microscopy, mathematical modelling and gene expression profiling of stationary phase *B. thailandensis* undergoing phage-antibiotic combination therapy as well as antibiotic or phage monotherapy.

## Results

### Genetic resistance to ΦBp-AMP1 phage in *B. thailandensis* is heterogeneous

We used well-mixed liquid cultures of stationary phase *B. thailandensis* (strain E264) with ΦBp-AMP1 at a multiplicity of infection (MOI) of 1 in lysogeny broth (LB) medium at 37 °C and performed colony forming unit (CFU) assays every 2 h over a 24 h period. For the first 14 h of incubation in the presence of phage, we did not observe neither a reduction nor an increase of the bacterial population, despite the presence of LB medium allowed regrowth of uninfected stationary phase *B. thailandensis* cultures (Figure 1A).

**Figure 1.**
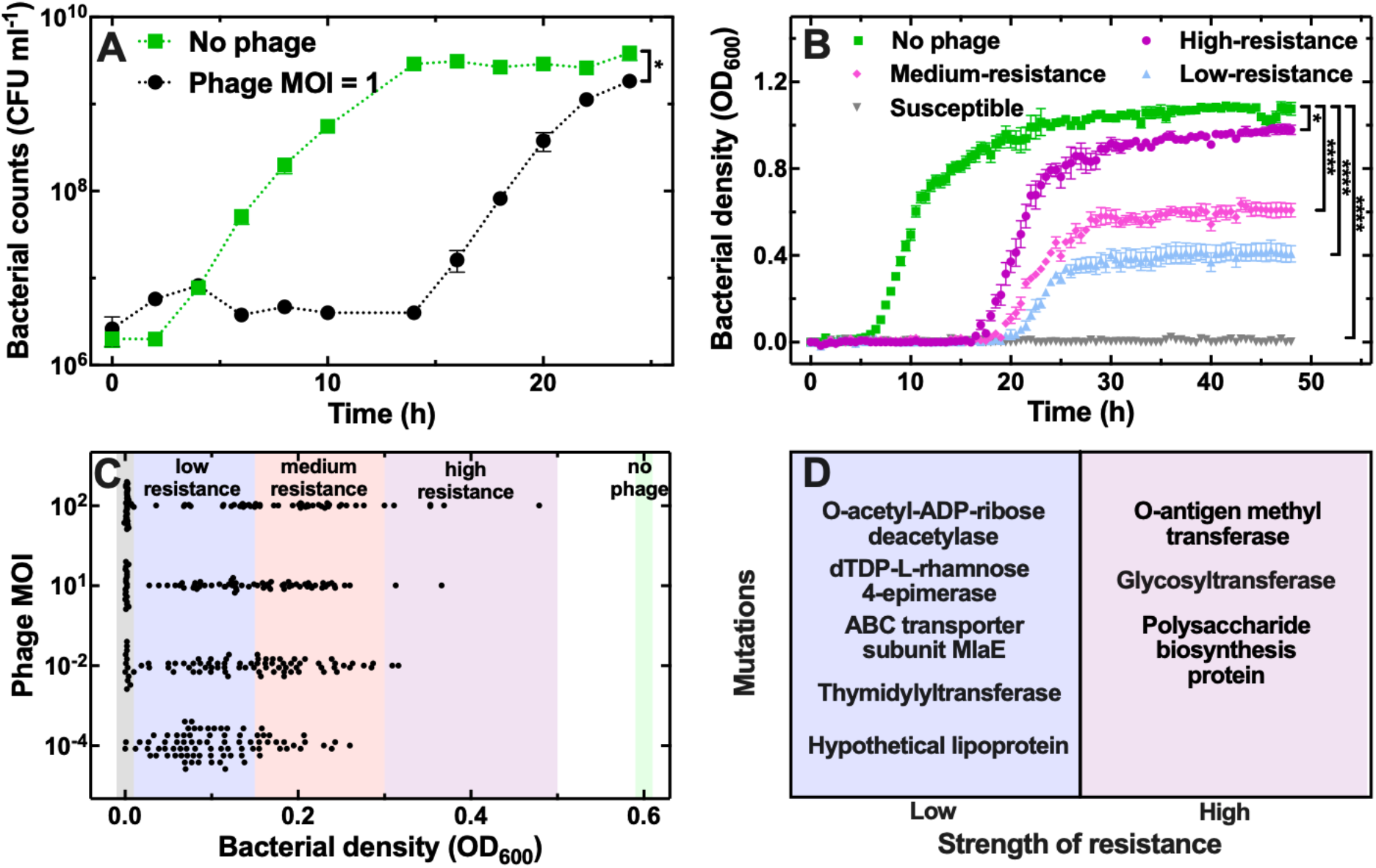
Heterogenous resistance to ΦBp-AMP1 in *B. thailandensis*. (A) Regrowth of stationary phase *B. thailandensis* populations in the presence of LB medium only (green squares) or together with phage at an MOI of 1 (black circles). Symbols and error bars are means and standard errors of the means of CFU measurements obtained from biological triplicates each containing technical triplicates. Very small error bars cannot be visualised due to overlap with the datapoints. Dotted lines are guides-for-the-eye. Corresponding phage counts are reported in Figure S1. (B) Regrowth of stationary phase *B. thailandensis* populations in the presence of LB medium only (1.05 < OD_600_ < 1.2 after 48 h, green squares) or together with phage at an MOI of 1 with different levels of bacterial resistance to phage emerging: high-resistance (0.9 < OD_600_ < 1.05 after 48 h, purple circles), medium-resistance (0.5 < OD_600_ < 0.7 after 48 h, magenta diamonds), low-resistance (0.3 < OD_600_ < 0.5 after 48 h, blue upward triangles), susceptible (0 < OD_600_ < 0.01 after 48 h, grey downward triangles). Symbols and error bars are means and standard errors of bacterial density values, measured in OD_600_, obtained from 84 technical replicates from biological triplicates. Very small error bars cannot be visualised due to overlap with the datapoints. * indicate a p-value < 0.05, **** indicate a p-value < 0.0001. (C). Bacterial density measurements after 24 h in the presence of phage at an MOI of 10^-4^, 10^-2^, 1 and 10^2^. Each black circle represents a bacterial density value performed on one of 84 technical micro-culture replicates from biological triplicates. The coloured vertical bands in the background indicate the range of OD_600_ values for each level of resistance after 24 h: high-resistance (0.3 < OD_600_ < 0.5, purple band), medium-resistance (0.15 < OD_600_ < 0.3, magenta band), low-resistance (0.01 < OD_600_ < 0.15, blue band), susceptibility (0 < OD_600_ < 0.01, grey band). (D) Unique mutations identified in five representative low- and high-resistance mutants. See Table S1 for further details.

The phage population instead increased within the first 4 h of co-incubation and reached a plateau after 6 h (Figure S1). The bacterial population started to increase after 14 h co-incubation with phage and after 24 h reached a plateau 2-fold lower compared to that measured in the absence of phage (Figure 1A). These data suggest that a bacterial subpopulation survives phage treatment, passes on phage immunity to its progeny and becomes the predominant genotype within the population; in contrast, a susceptible subpopulation becomes a minority due to phage infection but persists in the presence of phage since the phage population does not decline within our experimental time frame (Figure S1).

To test this hypothesis, we infected 84 different stationary phase bacterial micro-cultures and measured the bacterial density in the presence of phage and LB medium. We found that some bacterial micro-cultures contained only bacteria that were susceptible to phage and did not grow; whereas in other bacterial micro-cultures genetic resistance to phage emerged and the cultures started to regrow in the presence of LB medium, albeit with different onset times, slopes and saturations of growth (Figure 1B). Overall, the distribution of growth levels observed could be described by a hurdle-gamma distribution (Figure 1C and Figure S2), where the hurdle describes the cultures that are susceptible to phage, whereas the gamma describes the spread of growth in the cultures that are resistant to phage. These data suggest that *B*. *thailandensis* populations must contain a heterogeneous pool of mutants that are resistant to the phage.

Next, we set out to investigate whether the phage MOI has an impact on the emergence of a heterogeneous pool of genetically resistant mutants. We infected 84 stationary phase bacterial micro-cultures with four different phage MOIs and measured the bacterial density after 24 h. We found that the proportion of cultures susceptible to phage increased with the MOI; the fraction of cultures displaying low resistance to phage or high resistance to phage reduced and increased, respectively, with increasing phage MOI (Figure 1C and Figure S2). Moreover, all of the cultures grew when treated with ΦBp-AMP1 at an MOI of 1 at 25 °C (Figure S3), confirming that ΦBp-AMP1 features a temperature-dependent switch from the lytic to the lysogenic cycle.

To investigate the mechanisms underpinning resistance to phage, we harvested survivor populations from the experiments above and inoculated them either into 96-well plates or onto agar plates containing phage. We found that all survivors grew in the presence of ΦBp-AMP1 and LB medium, suggesting that the bacterial populations had become genetically resistant to phage. However, the colonies produced by these resistant mutants were smaller compared to the parental strain (Figure S4), suggesting that these mutations brought about a fitness cost ^24^. Therefore, we set out to understand which genetic mutations had emerged within the population during phage treatment. We sequenced the genome of five representative high-resistant and low-resistant survivor cultures. We found that low resistance was due to unique mutations in genes encoding the O-acetyl-ADP-ribose deacetylase, the dTDP-L-rhamnose-4-epimerase, the ABC transporter subunit MlaE, the thymidylyltransferase and a hypothetical lipoprotein (Figure 1D and Table S1). High resistance was instead due to unique mutations of three genes encoding the O-antigen methyl transferase, a glycosyltransferase and a polysaccharide biosynthesis protein. Moreover, we did not find lysogenic ΦBp-AMP1 in any of the genomes sequenced (Table S2), suggesting that stable lysogeny did not occur at 37°C. We found two common temperate bacteriophages of *Burkholderia* species, ΦE12-2 and ΦE125, in all sequenced *B. thailandensis* genomes (Table S2).

Taken together, these data suggest that ΦBp-AMP1 is a lytic phage of *B. thailandensis* at 37°C, that different levels of genetic resistance can emerge within putatively clonal populations of *B. thailandensis* depending on the phage MOI employed and that mutations of genes encoding membrane associated proteins confer high resistance to ΦBp-AMP1. Therefore, in order to make ΦBp-AMP1 effective in eradicating stationary phase *B. thailandensis* there is a need to use this phage in combination with clinically relevant antibiotics.

### ΦBp-AMP1 phage increases the growth inhibitory efficacy of quinolones, β-lactams and tetracyclines

We set out to investigate whether the use of ΦBp-AMP1 increases the efficacy of clinically relevant antibiotics in inhibiting the regrowth of stationary phase *B. thailandensis* when incubated in LB medium. We used four major antibiotic classes commonly employed for treatment of melioidosis, namely quinolones, β-lactams, tetracyclines and tetrahydrofolate synthesis inhibitors as well as antimicrobial agents that are not routinely employed to treat melioidosis, namely aminoglycosides, oxazolidinones, macrolides and glycopeptides (Table 1). We determined the fractional inhibitory concentration (FIC) index of each antibiotic against stationary phase *B. thailandensis* by dividing the minimum inhibitory concentration (MIC) derived from combination therapy with phage at an MOI of 1 by the MIC of antibiotic monotherapy. Therefore, the smaller the FIC index value measured, the higher is the increase in antibiotic efficacy of the combination therapy.

**Table 1.**
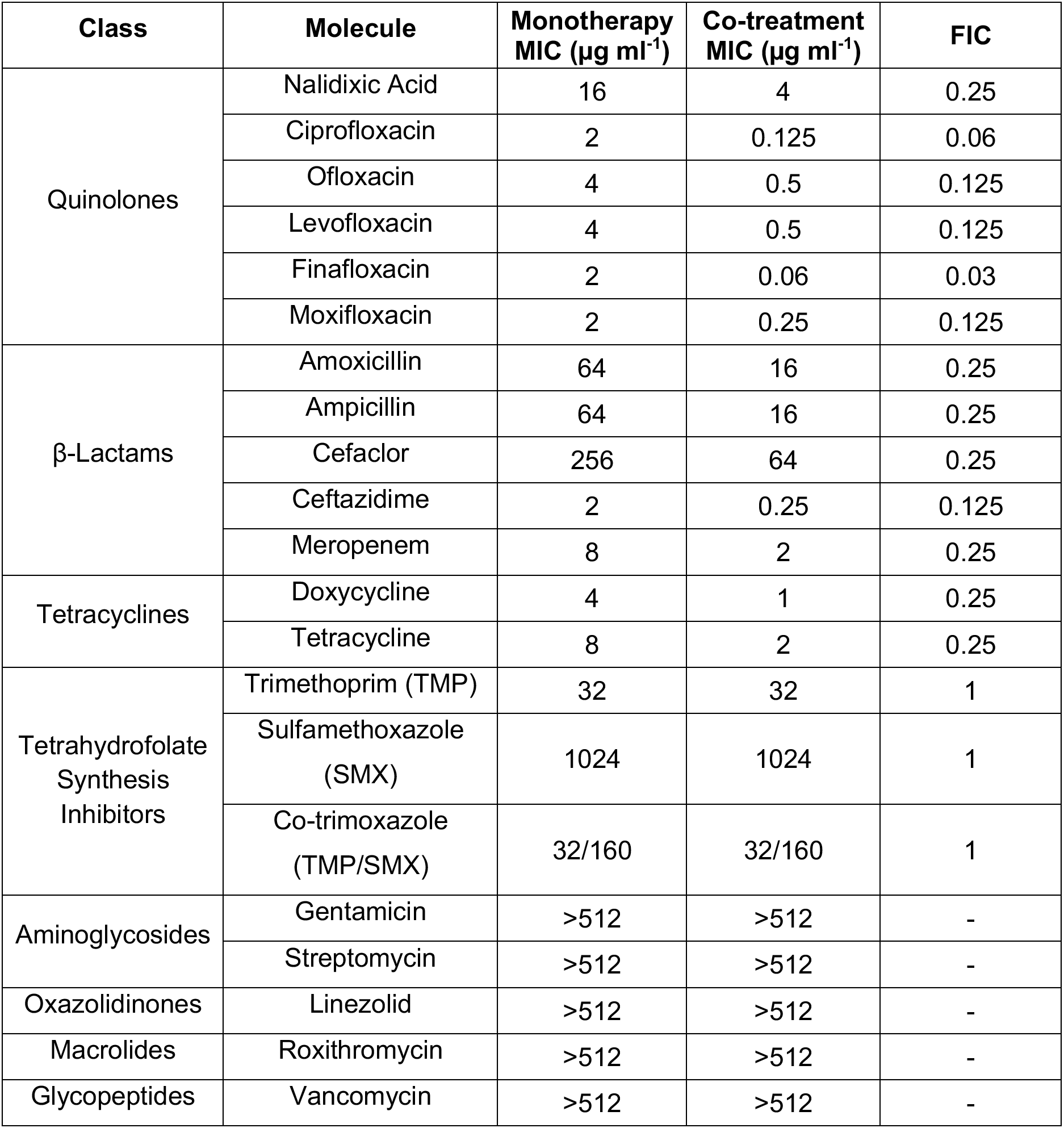
ΦBp-AMP1 phage increases the inhibitory efficacy of quinolones, β-lactams and tetracyclines. Antibiotic class, antibiotic molecule, minimum inhibitory concentration (MIC) measured against stationary phase *B. thailandensis* when used as monotherapy or in combination therapy with phage ΦBp-AMP1 at an MOI of 1 and the fractional inhibitory concentration (FIC) as the ratio of the MIC value obtained from the combination therapy over the MIC value obtained from monotherapy. The initial bacterial inoculum was 5×10^5^ cells mL^-1^, regrowth of stationary phase bacteria was measured as optical density after 24 h of monotherapy or combination therapy compared with regrowth of untreated stationary phase bacteria incubated in LB medium only. The MIC value after 24 h was determined as the antibiotic concentration value for which the bacterial growth was less than 10% the growth value measured for untreated bacteria. Each measurement was performed in biological triplicates each consisting of five technical replicates.

Remarkably, combination therapy increased the efficacy of a diverse range of molecules representative of all of the fluoroquinolone generations. Specifically, we measured an FIC index of 0.25 for nalidixic acid (1^st^ generation fluoroquinolone), 0.06 for ciprofloxacin (2^nd^ generation), 0.125 for levofloxacin (3^rd^ generation), 0.125 for moxifloxacin (4^th^ generation) and 0.03 for finafloxacin (5^th^ generation). Similarly, combination therapy increased the efficacy of representative β-lactams and tetracyclines against *B. thailandensis*; however, combination therapy did not increase the efficacy of tetrahydrofolate synthesis inhibitors (Table 1).

Inspired by these successful combination therapy findings, we set out to determine whether combination therapy with phage increased the efficacy of antibiotics that are not routinely employed to treat melioidosis. However, combination therapy with phage did not increase the efficacy of two aminoglycosides, an oxazolidinone, a macrolide or a glycopeptide (Table 1), antibiotics against which *B. thailandensis* is intrinsically resistant due to constitutively expressed efflux pumps and an atypical lipopolysaccharide structure ^2^.

Next, we investigated the bacterial population dynamics in the presence of phage and a sub-inhibitory concentration of levofloxacin, cefaclor or trimethoprim for which combination therapy with phage provided high, medium or no increase in efficacy, respectively (Table 1). Combination therapy with 0.25× MIC levofloxacin and phage at an MOI of 1 in LB medium did not allow for regrowth of stationary phase *B. thailandensis* within the 24 h experimental timeframe, whereas regrowth started after 5 h exposure to levofloxacin monotherapy and 16 h after exposure to phage monotherapy (Figure 2A).

**Figure 2.**
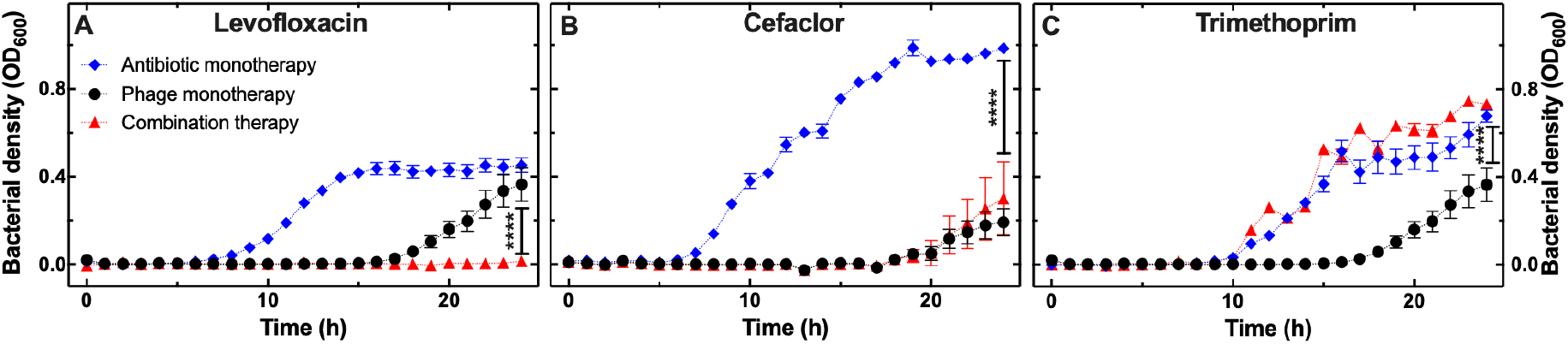
Bacterial population dynamics in the presence of ΦBp-AMP1 and sub-inhibitory antibiotic concentrations. Regrowth of stationary phase *B. thailandensis* populations in the presence of LB medium and either phage at an MOI of 1 (black circles), or 0.25× MIC of (A) levofloxacin, (B) cefaclor or (C) trimethoprim either as monotherapy (blue diamonds) or in combination therapy with phage at an MOI of 1 (red triangles). Symbols and error bars are means and standard errors of the means of bacterial density values, measured in OD_600_, obtained from biological triplicates each consisting of five technical replicates. Very small error bars cannot be visualised due to overlap with the datapoints. Dotted lines are guides-for-the-eye. **** indicate a p-value < 0.0001.

Following cefaclor monotherapy at 0.25× its MIC, the stationary phase *B. thailandensis* population started to regrow after 5 h, whereas following either phage monotherapy at an MOI of 1 or combination therapy with 0.25× MIC cefaclor and phage at an MOI of 1, the stationary phase *B. thailandensis* population started to regrow after 16 h and reached a plateau that was significantly lower compared to cefaclor monotherapy (Figure 2B). Following trimethoprim monotherapy at 0.25× its MIC, the stationary phase *B. thailandensis* population started to regrow after 9 h. Similarly, following combination therapy with 0.25× MIC trimethoprim and phage at an MOI of 1, the stationary phase *B. thailandensis* population started to regrow after 9 h and after 24 h reached a plateau that was significantly higher compared to that reached during phage monotherapy (red triangles and black circles in Figure 2C, respectively).

Taken together these data demonstrate that combination therapy with phage ΦBp-AMP1 increases the inhibitory efficacy of quinolones, β-lactams and tetracyclines, whilst combination therapy with tetrahydrofolate synthesis inhibitors decreases the inhibitory efficacy of ΦBp-AMP1.

### The interactions between phage ΦBp-AMP1 and antibiotics depend on the antibiotic but not on the phage concentration

Next, we set out to understand whether phage-antibiotic interactions in inhibiting bacterial growth depend on either the antibiotic or the phage concentration. Firstly, we measured the bacterial density of stationary phase *B. thailandensis* over time during exposure to LB medium and different antibiotic concentrations at a constant phage MOI of 1 (data points in Figure 3A, Figure 3F and Figure 3K). In order to estimate bacterial density and the associated uncertainty also for antibiotic concentrations that we did not investigate experimentally, we fitted these data to a statistical non-linear regression model (lines and bands in Figure 3A, Figure 3F and Figure 3K). This model also allowed us to infer the probability of additivism between the phage effect and the antibiotic effect on bacterial growth. We summarised our results in the form of interaction plots ^34^ reporting bacterial density values in the absence of phage or antibiotic, growth values following phage or antibiotic monotherapies and growth values following combination therapy.

**Figure 3.**
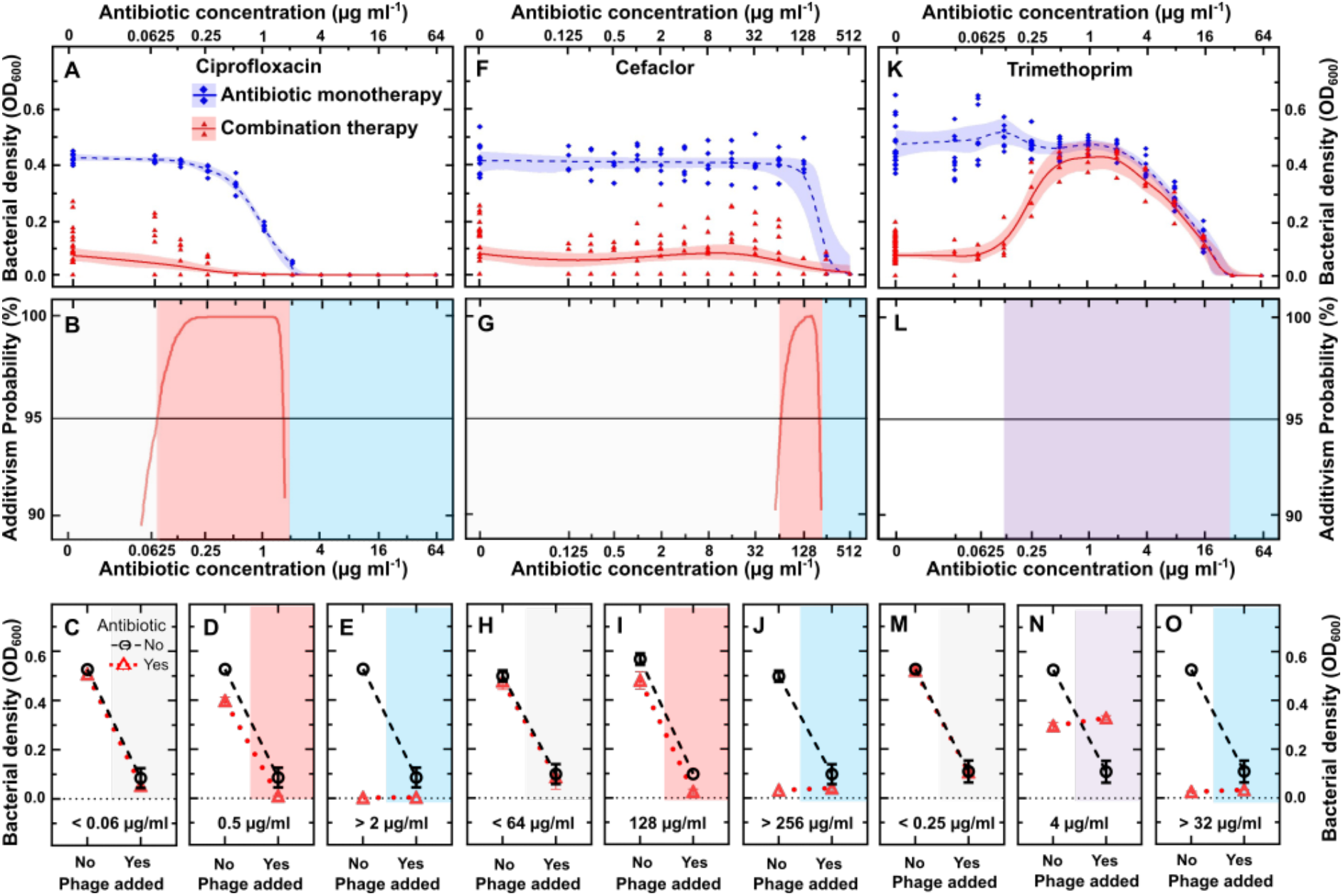
Interactions between ΦBp-AMP1 and antibiotics depend on the antibiotic concentration. Experimental data (symbols) and model predictions (lines and bands) describing the dependence of bacterial density on the concentration of (A) ciprofloxacin, (F) cefaclor or (K) trimethoprim in the absence (blue diamonds) and presence of phage ΦBp-AMP1 at an MOI of 1 (red triangles) after 24 h treatment. Each symbol represents the bacterial density measured in one of 15 technical replicates collated from biological triplicates. Some of the symbols overlap with each other. The lines and shaded areas are the medians, upper and lower quartiles, estimated by fitting our statistical non-linear regression model to our experimental data via Markov Chain Monte Carlo simulations. Corresponding predicted probability of an additive interaction between phage and (B) ciprofloxacin, (G) cefaclor and (L) trimethoprim (red lines) is shown only for antibiotic concentration ranges where the probability is higher than 90% (red shaded areas). Not shaded or blue shaded areas indicate antibiotic concentration ranges where the phage or the antibiotic dominate, respectively. Purple shaded areas indicate antagonism. Corresponding interaction plots at antibiotic concentrations selected from the ranges above for (C-E) ciprofloxacin, (H-J) cefaclor and (M-O) trimethoprim. Black circles connected by dashed lines show bacterial density values following control experiments and phage monotherapy, red triangles connected by dotted lines show bacterial density values following antibiotic monotherapy and combination therapy.

When ciprofloxacin was used above its MIC, we measured growth suppression following both monotherapy and combination therapy (Figure 3A), without additive effect (blue shaded area in Figure 3B and Figure 3E) in line with previous reports ^34^. In contrast, when ciprofloxacin was used at sub-inhibitory concentrations, the model predicted an interacting region: successful growth suppression was achieved only in the case of combination therapy (red lines in Figure 3A), with a degree of confidence of >95% in predicted additive effect at ciprofloxacin concentrations between 0.06 and 2 µg ml^-1^ (red shaded area in Figure 3B and Figure 3D). This additive concentration range was broad for ciprofloxacin, finafloxacin and moxifloxacin and narrower for nalidixic acid, ofloxacin and levofloxacin (Table S3). Finally, we did not record an interaction effect between phage and ciprofloxacin at low concentrations (white shaded area in Figure 3B and Figure 3C) in line with previous reports ^34^.

We recorded similar concentration-dependent interactions between phage and cefaclor (Figure 3F-J); however, for this antibiotic the additive effect was limited to the antibiotic range 64-256 µg mL^-1^, i.e. a 4-fold additive concentration range (Figure 3G). In addition, ceftazidime, meropenem, doxycycline, tetracycline, amoxicillin and ampicillin displayed a narrow additive concentration range (Table S3).

In contrast, we found an antagonistic effect when phage was used in combination with trimethoprim concentrations in the range of 0.125-32 µg mL^-1^ with the line connecting the growth value following antibiotic therapy and combination therapy (red, dotted line in Figure 3N) having a less negative slope than the line connecting the growth value following control experiments and phage monotherapy (black, dashed line in Figure 3N). A similarly extended antagonistic range was recorded when phage was used in combination with either sulfamethoxazole or co-trimoxazole.

Secondly, we measured the bacterial density of stationary phase *B. thailandensis* over time during incubation in LB medium while simultaneously varying the initial antibiotic concentration and phage MOI. We used ciprofloxacin, ampicillin and trimethoprim as representative molecules displaying a broad additive concentration range, a narrow additive concentration range and antagonism, respectively, in combination with phage at an initial MOI of either 10^-4^, 10^-2^, 10^0^ or 10^2^. We found that growth inhibition was significantly extended below the MIC of ciprofloxacin or ampicillin even at a phage MOI of 10^-4^ (Figure 4A and Figure 4B), suggesting that a modest phage concentration is sufficient for increasing ciprofloxacin and ampicillin inhibitory efficacy against *B. thailandensis*.

**Figure 4.**
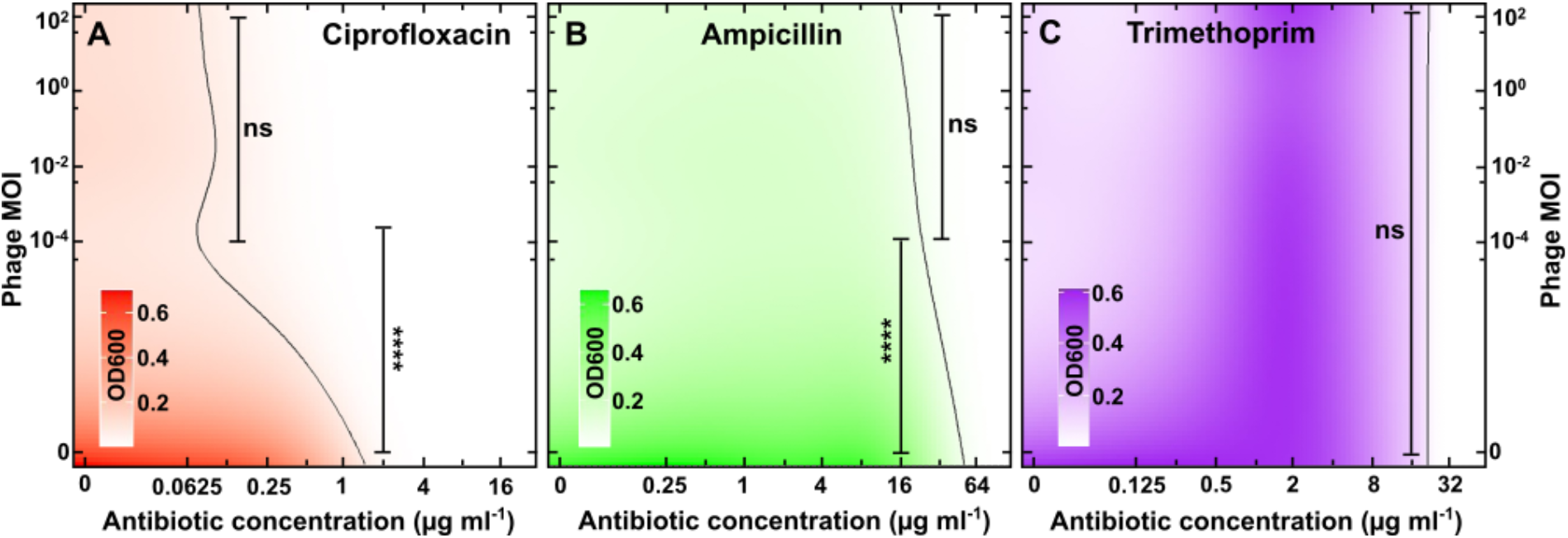
Phage-antibiotic interactions are not affected by the phage concentration. Heatmaps of *B. thailandensis* density (measured in OD_600_) after 24 h treatment with different initial phage MOIs and different concentrations of (A) ciprofloxacin, (B) ampicillin or (C) trimethoprim. Heatmaps were obtained via hierarchical Bayesian statistical modelling fitted to our experimental data (measured only at the phage and antibiotic concentrations indicated on the x- and y-axes). The vertical black lines are predictions of all antibiotic-phage combinations that permit bacterial density values that are lower than 10% of the bacterial density values obtained for bacteria growing in LB medium only. ns indicates no statistical significance, **** indicates a p-value < 0.0001.

In contrast, growth inhibition was not extended below the MIC of trimethoprim at any of the phage MOIs tested (Figure 4C). Moreover, in the presence of phage *B. thailandensis* growth was maximal for trimethoprim concentrations in the range 1 - 8 µg mL^-1^ and decreased at lower trimethoprim concentrations (Figure 4C). Therefore, these data confirm the hypothesis above that sub-inhibitory concentrations of trimethoprim antagonize with phage efficacy to inhibit bacterial growth. Taken together, these data demonstrate that the interaction between phage ΦBp-AMP1 and antibiotics strongly depends on the antibiotic concentration and mode of action but not on the initial phage MOI.

### Synergistic bactericidal effect of phage and ciprofloxacin in combination therapy

We next set out to quantify the bactericidal efficacy of phage-antibiotic combination therapy. We exposed stationary phase *B. thailandensis* to a range of concentrations of ciprofloxacin and phage at an MOI of 1 for 24 h, we quantified bactericidal efficacy by measuring survivors via CFU assays and calculated the survivor fold reduction compared to untreated bacteria (i.e. the higher the fold reduction in Figure 5A the stronger the bactericidal effect). Combined with phage, sub-MIC concentrations of ciprofloxacin allowed for a survivor fold reduction that was between 3 and 120 times greater compared to ciprofloxacin monotherapy (red and blue bars in Figure 5A, respectively, p-value < 0.0001 for all pair-wise comparisons); thus, suggesting a synergistic bactericidal effect between phage and ciprofloxacin, considering that phage monotherapy provided only a 2-fold survivor reduction (Figure 5A). Moreover, at supra-MIC concentrations, combination therapy achieved a survivor fold reduction that was between 570 and 2400 times greater compared to ciprofloxacin monotherapy (p-value < 0.0001 for all pair-wise comparisons a part from 8× MIC). Notably, the minimum bactericidal concentration that provided complete eradication of stationary phase *B. thailandensis* (horizontal band in Figure 5A) was 8× MIC for ciprofloxacin monotherapy and 4× MIC for phage-ciprofloxacin

**Figure 5.**
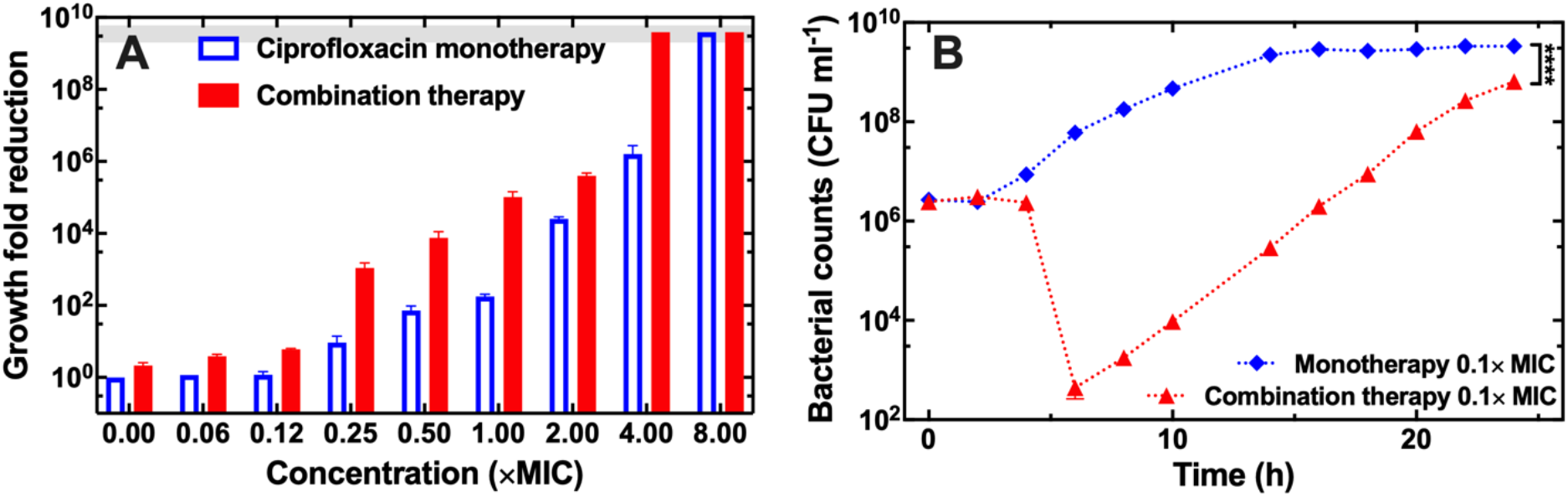
Synergistic bactericidal effect of phage and ciprofloxacin combination therapy. (A) Dependence of the ratio between the number of bacteria viable after 24 h incubation in LB medium and the number of bacteria viable 24 h ciprofloxacin monotherapy (blue bars) or phage-ciprofloxacin combination therapy (red bars) on the concentration of ciprofloxacin employed. In all cases the starting inoculum was stationary phase *B. thailandensis* at a concentration of 5×10^5^ CFU mL^-1^ and bacteria were counted at 24 h via CFU assays. The grey horizontal band represents a survival fold reduction that corresponds to the complete eradication of the bacterial population, i.e. colony counts below limit of detection of 10 CFU mL^-1^. Pair-wise t-tests revealed that combination therapy yielded a significantly (p-value < 0.0001) higher growth reduction at all ciprofloxacin concentrations tested compared to ciprofloxacin monotherapy, a part from 8× MIC. (B) Temporal dependence of bacterial counts following 0.125× MIC ciprofloxacin monotherapy (blue diamonds) or combination therapy with 0.125× MIC ciprofloxacin and phage at an MOI of 1 (red triangles). Symbols and error bars are means and standard errors of the mean of biological triplicates each containing technical triplicates. Very small error bars cannot be visualised due to overlap with the datapoints. Dotted lines are guides-for-the-eye.

Next, we set out to measure and contrast the dynamics of the bactericidal effect of mono- and combination therapy at a sub-MIC antibiotic concentration. We treated stationary phase *B. thailandensis* with either ciprofloxacin at 0.125× its MIC or a combination of phage at an MOI of 1 and ciprofloxacin at 0.125× its MIC and measured the number of survivors at regular time points via CFU assays (Figure 5B). Following ciprofloxacin monotherapy, the bacterial population was not affected by antibiotic exposure but started expanding after 2h of treatment and reached stationary phase after 14h of treatment (blue diamonds in Figure 5B). In contrast, following combination therapy the bacterial population reduced between 4 h and 6 h and reached a minimum that was >5-log lower compared to the initial bacterial inoculum (red triangles in Figure 5B). The surviving bacterial population started to increase after 6 h of treatment due to the emergence of phage resistance (Figure S5) and reached a maximum level at 24 h that was 1-log lower than bacteria treated with ciprofloxacin monotherapy (red triangles and blue diamonds in Figure 5B, respectively). Taken together these data confirm that the use of ΦBp-AMP1 allows a reduction in the concentration of antibiotic required to both inhibit growth of or kill stationary phase *B. thailandensis*.

### Synergistic bactericidal interactions between phage and antibiotics are underpinned by phage-induced downregulation of efflux in *B. thailandensis*

Next, we set out to discover the molecular mechanisms underpinning the newly found synergistic bactericidal interaction between ΦBp-AMP1 and ciprofloxacin. We treated stationary phase *B. thailandensis* for 4 h either with 0.125× MIC ciprofloxacin monotherapy or phage monotherapy at an MOI of 1 or combination therapy with both 0.125× MIC ciprofloxacin and phage at an MOI of 1. We chose 4 h-long treatments because these treatments returned similar survivor numbers before the > 5-log reduction in survivor numbers measured for combination therapy at the 6 h time point (Figure 5B). We extracted bacterial and phage RNA from biological triplicates of each condition and performed a global comparative transcriptomic analysis ^39^ among the transcriptomes obtained from these three different conditions and with respect to stationary phase *B. thailandensis* incubated in LB medium for 4 h (Supplementary files 1-3). Using principal component analysis, we found that bacterial transcriptome replicates from each condition clustered together and were well separated from replicates from different conditions (Figure S6A). The only exception were the bacterial transcriptomes harvested from cultures treated with phage mono- and combination therapy (black circles and red triangles in Figure S6A) that largely overlapped, suggesting that, at the concentrations employed, phage had a greater impact than ciprofloxacin on the bacterial transcriptomes. Moreover, the phage transcriptomes harvested from cultures treated with mono- and combination therapies also largely overlapped according to our principal component analysis (Figure S6B).

Gene ontology enrichment analysis ^40^ of differentially expressed genes revealed that all treatments investigated resulted in the downregulation of locomotion processes, in accordance with a recent study using *P. aeruginosa* ^41^, as well as cell localization, cell projection, organelle organization and response to external stimuli processes (Figure 6A, 6B and Supplementary files 4-6). Ciprofloxacin monotherapy also caused the downregulation of biological processes involved in cell communication, as well as the upregulation of RNA, catabolic, metabolic and ribosomal processes (Figure 6A and Supplementary file 4).

**Figure 6.**
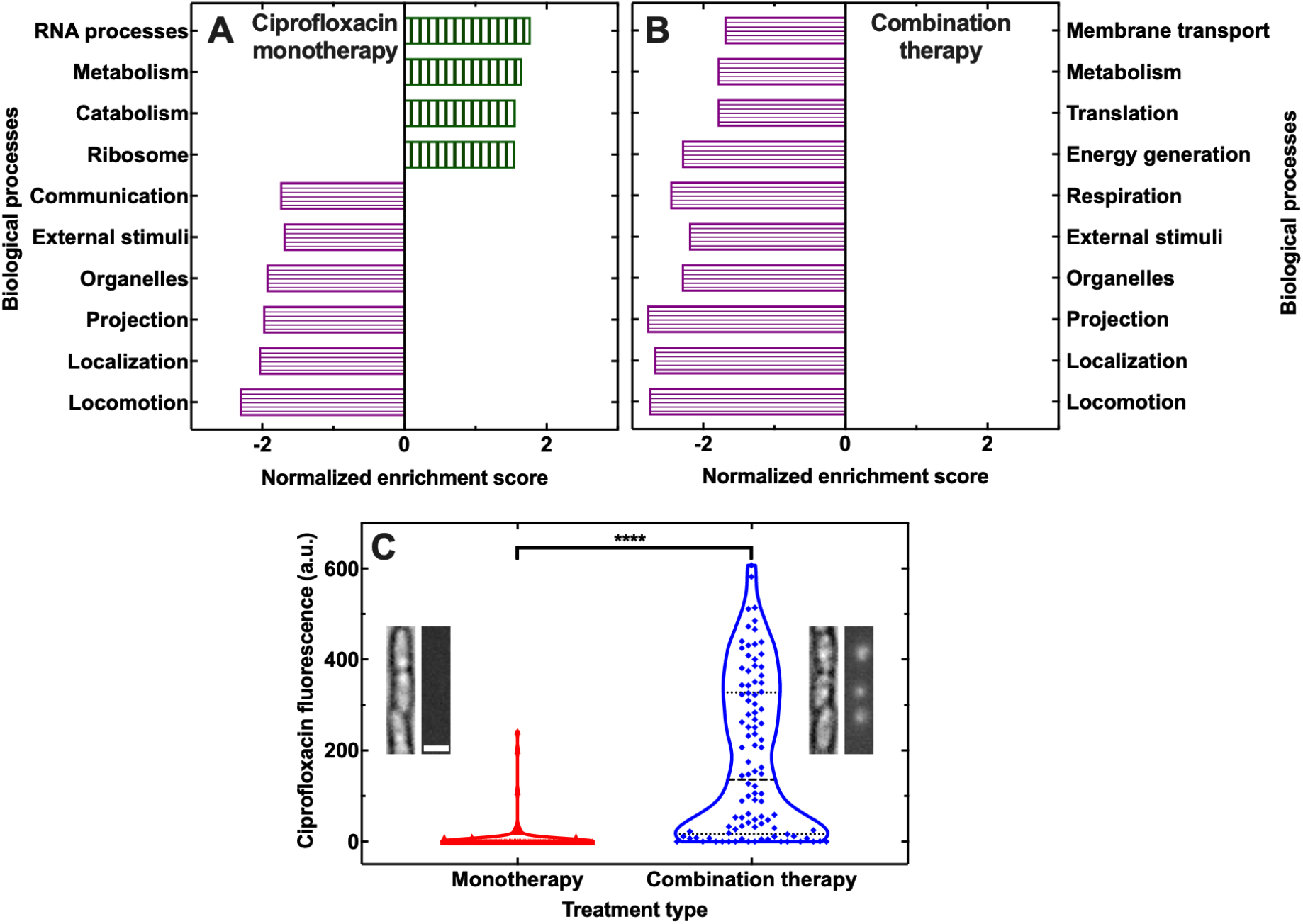
Synergistic bactericidal interactions between phage and ciprofloxacin are underpinned by phage-induced downregulation of membrane transport in *B. thailandensis*. Major biological processes that are significantly enriched and either up- or downregulated (vertically and horizontally patterned bars, respectively) in stationary phase *B. thailandensis* after 4 h of (A) monotherapy with ciprofloxacin at 0.125× MIC or (B) combination therapy with ciprofloxacin at 0.125× MIC and phage at an MOI of 1 with respect to untreated *B. thailandensis* incubated for 4 h in LB medium only. Corresponding differential gene expression analyses and gene ontology enrichment analyses are reported in Supplementary files 1-3 and 4-6, respectively. (C) Distribution of values of fluorescence of ciprofloxacin-NBD accumulating in N = 100 individual stationary phase *B. thailandensis* that had been incubated for 4 h in either ciprofloxacin-NBD only or ciprofloxacin-NBD and phage (red triangles and blue diamonds, respectively). Ciprofloxacin-NBD was introduced in the microfluidic device at t = 0 at a concentration of 32 µg ml^-1^, phage was introduced at t = 0 at a concentration of 10^8^ PFU ml^-1^. Dashed lines indicate the median of each distribution, dotted lines indicate the quartiles of each distribution. **** indicate a p-value < 0.0001.

In contrast, both phage monotherapy and combination therapy caused the downregulation of metabolic, membrane transport, translational, energy generation and respiration processes (Figure 6B and Supplementary files 5 and 6). Specifically, genes encoding the major efflux pump BpeEF-OprC were significantly downregulated along with functional subunits of the F_0_F_1_-type ATP synthases, of the cytochrome O oxidase, of the NADH-quinone oxidoreductase complex, of lipopolysaccharide transport and of secretion systems (Supplementary file 2).

Combination therapy drove a global differential gene regulation that was very similar to that caused by phage monotherapy and gene ontology enrichment analysis did not return any enriched functional category for this comparison. Moreover, the expression of phage encoded genes was also unaffected by the presence of ciprofloxacin (Supplementary file 7) and gene ontology enrichment analysis did not return any enriched functional category for this comparison either.

Next, we set out to test the hypothesis that phage-induced downregulation of efflux processes led to increased intracellular accumulation of ciprofloxacin during phage-ciprofloxacin combination therapy. We used our recently reported microfluidics-based time-lapse microscopy platform ^42,43^ and a fluorescent derivative of ciprofloxacin (i.e. ciprofloxacin-nitrobenzoxadiazole, henceforth ciprofloxacin-NBD) to measure intracellular accumulation of ciprofloxacin in individual stationary phase *B. thailandensis* cells. In accordance with our hypothesis, we found that the distribution of ciprofloxacin-NBD fluorescence values after 4 h phage-ciprofloxacin-NBD combination therapy was significantly higher than the distribution of ciprofloxacin-NBD fluorescence values after 4 h ciprofloxacin-NBD monotherapy (Figure 6C).

Moreover, using this platform we did not find evidence of either cell filamentation or a significant difference in cell size between stationary phase *B. thailandensis* incubated in LB growth medium only (Figure S7 A-E), or during phage monotherapy (Figure S7 F-J) or during combination therapy with phage at an MOI of 1 and ciprofloxacin at 0.125× MIC (Figure S7 K-O). We found that stationary phase *B. thailandensis* infected with phage in the presence of sub-inhibitory concentrations of ampicillin or ciprofloxacin produced less phage particles than in the absence of antibiotics in planktonic cultures and plaques of similar size on agar cultures (Figure S1 and S8, respectively). Sub-inhibitory concentrations of trimethoprim also led to smaller plaques with phage propagation starting significantly later in the presence of trimethoprim (Figure S1 and S8). We also did not observe plaque formation on LB agar plates in the presence of sub-inhibitory concentrations of ciprofloxacin in the absence of phage ΦBp-AMP1. Moreover, in our phage transcriptome analysis we found evidence of expression of structural phage proteins, but we did not find evidence of expression of genes encoding excisionases, i.e. proteins required for the excision of dormant phage from within the hosts genome ^44^, neither during phage mono- nor combination therapy (Supplementary file 7). However, it is conceivable that one or more of the 11 hypothetical proteins expressed by ΦBp-AMP1 (Supplementary file 7) could perform excisionase activity.

Taken together these data demonstrate that the observed phage-antibiotic interactions are not due to cell filamentation, increase phage particle production or phage induction in the presence of antibiotics but are instead underpinned by phage-induced downregulation of membrane transport and energetic processes that are involved in the efflux of ciprofloxacin leading to higher intracellular ciprofloxacin accumulation in the presence of phage.

## Discussion

### Emergence of resistance to phage

Phage and bacteria are engaged in a constant arms race leading to the evolution of a multitude of non-mutually exclusive antiphage defence mechanisms, including the well-understood phage receptor alteration, restriction-modification, abortive infection, CRISPR-Cas systems, as well as new defence systems ^45,46^. It is well established that increased phage virulence selects for the evolution of host resistance if the costs associated with resistance are outweighed by the benefits of the capability to avoid infection ^47,48^. Accordingly, we found that the level of resistance to phage increased with the strength of phage predation. Interestingly, even within a putatively clonal *B. thailandensis* population we found evidence of emergence of different levels of resistance due to mutations in eight different genes. Three of these genes encoded glycosyltransferase and O-antigen synthesis, that are strongly linked with the LPS ^49^, and capsular polysaccharides. Our data therefore suggest that the LPS or capsular polysaccharides could be the receptors for phage ΦBp-AMP1. This hypothesis is further corroborated by our global comparative transcriptomics analysis demonstrating the downregulation of LPS assembly associated genes following exposure to ΦBp-AMP1. Indeed, the LPS is a well-known receptor for many different bacteriophages and mutations of LPS confer resistance to phage in a variety of bacteria ^50–53^, whereas capsular polysaccharides have recently been shown to serve as primary receptors of *E. coli* phage ^54,55^. Moreover, it has been reported that *B. pseudomallei* displays various colony morphotypes based on O-antigen variations ^56^. Accordingly, we have demonstrated that phage-resistant bacteria form smaller colonies compared with the parental strain while displaying mutations in genes associated with O-antigen synthesis.

### Phage-antibiotic interactions

The interactions between two or more antimicrobials as components of combination therapies are broadly classified in three main types: additive (the sum of the effect of each component), synergistic (a larger-than-additive effect) and antagonistic (a smaller-than-additive effect ^13,57,58^). In the context of phage-antibiotic therapy instead, phage-antibiotic synergy has been defined as the stimulation of phage replication when bacteria are treated with sub-inhibitory concentrations of antibiotics ^28,29,59^. Considering that in accordance with a recent report ^60^ we did not find stimulation of phage replication in the presence of antibiotics, we chose to use the more broadly accepted definitions of additive, synergistic and antagonistic interactions, introduced above ^13,57,58^. Specifically, by using statistical analysis on interaction plots we found an additive effect between the phage and most of the tested molecules from the quinolone, β-lactam and tetracycline antibiotic classes, antagonism between the phage and tetrahydrofolate synthesis inhibitors and bactericidal synergy between the phage and the quinolone ciprofloxacin. These effects were comparable or more efficient in suppressing bacterial growth than previously reported synergistic phage-antibiotic effects ^28,33,34^. For example, ceftazidime efficacy increased by a factor of 2 compared to ceftazidime monotherapy when used in combination with the podovirus vB_BpP_HN01 ^61^ or the myovirus KS12 ^9^ against *B. pseudomallei* or *B. cenocepacia*, respectively; we measured an 8-fold increase in ceftazidime efficacy against *B. thailandensis* in the presence of phage ΦBp-AMP1. Using the temperate phage HK97 in combination with ciprofloxacin, a previous study has reported bactericidal synergy against *E. coli* K12, with complete eradication being achieved at 0.5× MIC ciprofloxacin ^32^; we measured a comparatively weaker bactericidal synergy between ΦBp-AMP1 and ciprofloxacin, with complete eradication being achieved at 4× MIC ciprofloxacin. However, it is worth noting that stationary phase *B. thailandensis* is significantly more resistant than exponential phase *E. coli* K12 with ciprofloxacin MIC values of 2 and 0.2 µg/ml, respectively.

### Factors affecting phage-antibiotic interactions

The dependence of phage-antibiotic interactions on the antibiotic class is well established ^28,29,34,59,60,62^, however, the mechanisms underpinning this dependence remain largely unknown. For example, the myovirus KS12 broadly synergised with quinolones, β-lactams and tetracycline when used to inhibit growth of *B. cenocepacia* but antagonised with aminoglycosides ^29^. The myovirus ΦHP3 displayed a synergistic effect with ceftazidime, an additive effect with kanamycin and an antagonistic effect with chloramphenicol when used to inhibit growth of *E. coli* ^34^. The phage PYO^SA^ antagonised tetracycline, azithromycin, and linezolid but synergised with daptomycin, vancomycin and kanamycin when used to inhibit growth of *S. aureus* ^33^. Here we advance this understanding by showing that even molecules within the same class can display a dramatically different extent and range of additive interactions with the same phage and that when used with phage the same molecule can simultaneously display an additive effect in inhibiting bacterial growth and a synergistic effect in killing bacteria.

The outcome of phage-antibiotic therapy is often contradictory because of a lack of systematic analysis of interactions between phage and antibiotics: these interactions are often studied with only one or two concentrations of the antimicrobials, which are wholly insufficient in predicting combinatorial concentrations that are effective during treatment ^20^. By assessing bacterial growth when exposed to multiple orders of magnitude of antibiotic concentrations and phage titers, we discovered that phage-antibiotic interactions strongly depend on the antibiotic concentration employed, but surprisingly do not vary with phage titer, a key difference from previous findings ^34^. These newly discovered dependences should be taken into account when designing rational phage-antibiotic therapy ^20,21^. In fact, our data suggest that it might be relatively straightforward to hit a suitable ΦBp-AMP1 phage titer in an *in vivo* setting, where phage are broadly tolerated ^63^ but their pharmacokinetic and pharmacodynamic parameters are less known compared to antibiotics ^64^.

### Mechanistic understanding of phage-antibiotic interactions

The additive effects between phage and antibiotics are stronger and broader in our experiments compared to previous reports ^29,59,61^, possibly due to differences at the phage level, i.e., a podovirus vs a myovirus, at the bacterial strain and physiology level, i.e., stationary phase *B. thailandensis* vs exponential phase *B. cenocepacia*, or due to a different mechanism of interaction between phage and antibiotics. Indeed, KS12 and a variety of other phage displayed an increase in plaque size and phage titer in the presence of sub-inhibitory concentrations of antibiotics ^26,28,31,59^, possibly caused by the acceleration of phage assembly and cell lysis due to cell filamentation in the presence of antibiotics ^28,29,34,65^. In contrast, we did not find evidence neither of cell filamentation nor of an increase in plaque size and phage titer in the presence of sub-inhibitory concentrations of quinolones, β-lactams or tetracyclines. Reduced phage titer in the presence of sub-inhibitory concentrations of trimethoprim could instead explain the observed antagonism between ΦBp-AMP1 and trimethoprim. Moreover, temperate phage activity is known to enhance antibiotic efficacy through depletion of lysogens ^32^. Although we found evidence of prophage ΦE125 and ΦE12-2 ^85,86^, it is unlikely that depletion of ΦE125 or ΦE12-2 lysogens plays a role in our experiments since we did not find evidence of plaque formation in the presence of ciprofloxacin, that is a lysogen activating antibiotic ^32^, in the absence of ΦBp-AMP1.

Based on our global gene expression analysis, we hypothesized that antibiotics are more effective in inhibiting *B. thailandensis* growth in the presence of ΦBp-AMP1 due to phage-induced down-regulation of antibiotic efflux out of the cell. Indeed, the multi-drug efflux pump BpeEF-OprC ^8^ was downregulated in the presence of phage and it is known that deletion of BpeEF-OprC causes increased susceptibility to quinolones and β-lactams ^9^. Moreover, we detected a strong downregulation of genes associated with aerobic respiration and transmembrane transport of protons, effectively reducing the availability of ATP for active transport of substrates as well as the proton motive force. A reduction in proton motive force levels leads to reduced antibiotic efflux ^66–68^, and in the long-term selection for mutations in drug efflux components ^69^. In accordance with our hypothesis, we found that a fluorescent derivative of ciprofloxacin accumulates in individual *B. thailandensis* cells at significantly higher levels in the presence of ΦBp-AMP1 compared with in its absence. Noteworthy, previous studies using phage that have the efflux component TolC as a receptor, demonstrated that emergence of phage resistance in bacteria via *tolC* mutations led to increased antibiotic susceptibility ^52,70^. Finally, it is conceivable that the phage-antibiotic interactions we observed are also due to phage and antibiotics targeting cells in different metabolic states as recently hypothesised ^60^.

In summary, the interactions between phage and bacteria are multifaceted and complex due to a billion-year long arms race between both entities ^71–75^ and are further modified when a second selective pressure is imposed, such as the presence of antibiotic compounds secreted by other microbes in the environment ^76^. Understanding such interactions might hold the key for successful antimicrobial therapy and to overcome the current antimicrobial resistance crisis. Considering that stationary phase bacteria are traditionally refractory to antibiotics, especially in spatial structures such as biofilms where antibiotic diffusion is further hindered ^42^, our data offer a potential route for their eradication by combining low doses of clinically relevant antibiotics with low doses of phage, that should be easily obtainable *in vivo* thanks to the self-propagating nature of phage.

## Materials and Methods

### Key Resources Table

| REAGENT or RESOURCE | Source | Identifier |
| --- | --- | --- |
| **Bacterial and phage strains** |  |  |
| *B. thailandensis* E264 | (Kim et al., 2005) | RefSeq: NC_007650/51 |
| ΦBp-AMP1 | (Gatedee et al., 2011) | RefSeq: NC_047743 |
| **Chemicals** |  |  |
| Amoxicillin | Sigma-Aldrich | Cat# A8523 |
| Ampicillin | Sigma-Aldrich | Cat# A9518 |
| Cefaclor | Sigma-Aldrich | Cat# C6895 |
| Ceftazidime | Melford | Cat# C59200 |
| Ciprofloxacin | Sigma-Aldrich | Cat# 17850 |
| Doxycycline | Sigma-Aldrich | Cat# D9891 |
| Finafloxacin | Sigma-Aldrich | Cat# SML2134 |
| Levofloxacin | Sigma-Aldrich | Cat# 40922 |
| Linezolid | Sigma-Aldrich | Cat# O9409 |
| Meropenem | Sigma-Aldrich | Cat# M2574 |
| Moxifloxacin | Sigma-Aldrich | Cat# PHR1542 |
| Nalidixic acid | Sigma-Aldrich | Cat# N8878 |
| Ofloxacin | Sigma-Aldrich | Cat# O8757 |
| Roxithromycin | Sigma-Aldrich | Cat# R4393 |
| Sulfamethoxazole | Sigma-Aldrich | Cat# S7507 |
| Tetracycline | Sigma-Aldrich | Cat# 87128 |
| Trimethoprim | Sigma-Aldrich | Cat# T7883 |
| Vancomycin | Sigma-Aldrich | Cat# 94747 |
| **Software and algorithms** |  |  |
| R version 3.4.1 | (R Core Team, 2017) | https://cran.r-project.org |
| brms R package | (Bürkner, 2017) | https://cran.r-project.org |
| breseq | (Deathrager and Barrick, 2012) | https://barrickslab.org |
| ImageJ | (Schneider et al., 2012) | https://imagej.nih.gov |
| GraphPad Prism 8.3.0 | GraphPad | https://www.graphpad.com/ |
| PHASTER | University of Alberta, Canada | https://phaster.ca |
| **Other** |  |  |
| CLARIOstar plate reader | BMG Labtech |  |
| WPA cell density meter | Scientific Laboratory Supplies | Cat# CO8000 |
| Sartorius Minisart™ 0.2 µm | Fisher Scientific | Cat# 10730792 |
| RNeasy Mini Kit (50) | QIAGEN | Cat# 74104 |
| Qubit 1.0 Fluorometer | Invitrogen | Cat# Q32857 |
| IX73 | Olympus, Tokyo, Japan |  |
| UPLSAPO60XW | Olympus, Tokyo, Japan |  |
| Flow Unit S | Fluigent |  |
| MFCs-4C | Fluigent |  |
| Zyla 4.2 | Andor |  |

### Bacterial and bacteriophage strains

*B. thailandensis* E264 strain and the temperature dependent, lytic bacteriophage ΦBp-AMP1 were obtained from the Department of Immunology, Faculty of Medicine Siriraj Hospital, Mahidol University.

### Bacterial culturing

*B. thailandensis* was stored at -80 °C and streaked on Lysogeny broth (LB, 10 g/L Tryptone, 5 g/L Yeast extract, 10 g/L NaCl, Melford) agar plates (10 g/L, 1.5% Agar) every two weeks. Overnight cultures were setup by inoculating a single *B. thailandensis* colony from a plate into flasks containing 50 mL of LB broth and were grown for 17 h at 37 °C on shaking platforms set at 200 rpm.

### Propagation and titration of phage

Phage propagation was carried out as previously reported ^77^. Briefly, overnight cultures of *B. thailandensis* E264 were diluted 1000× in 50 mL LB medium to obtain a bacterial concentration of approximately 2×10^6^ CFU mL^-1^. These sub-cultures were then incubated for 4 h at 37 °C and 200 rpm, allowing them to reach early exponential phase (∼8×10^6^ CFU mL^-1^). Then, these sub-cultures were infected with ΦBp-AMP1 at an MOI = 0.01 and incubated for 17 h at 37 °C and 200 rpm. On the following day, cells were pelleted by centrifugation for 40 minutes at 3000 *g* and the phage-containing supernatant was filtered twice (Sartorius Minisart^TM^ 0.2 µm) to obtain a phage stock. The phage concentration within this stock was then determined via the double agar overlay technique ^76^. Briefly, LB top agar (10 g/L, 0.5% Agar) was melted in a microwave and allowed to cool to below 40 °C. An aliquot of an overnight *B. thailandensis* E264 culture was added to the melted top agar at a volume: volume concentration of 1:50. LB agar plates were subsequently coated in a thin layer of the top agar above creating a continuous bacterial lawn. In parallel, a 10-fold dilution series of the phage stock was prepared in LB medium. Phage was then titred using the standard double agar overlay technique ^103^. All plates were then incubated at 37 °C for 17 h after which the phage induced plaques in the bacterial lawn were counted and the plaque forming units (PFU) per mL^-1^ of the phage stock was calculated. Phage stocks were stored at 4 °C for a maximum of two weeks before a new propagation was performed. Prior to each use, the phage concentration within the phage stock was determined via the double agar overlay technique above.

### Bacterial growth in the presence of phage

In order to measure the growth of bacteria in the presence of phage, we used two independent approaches. Firstly, we measured the concentration of bacteria during phage infection over time using colony forming unit (CFU) assays ^78^. Briefly, three overnight cultures of *B. thailandensis* were diluted in 50 mL LB medium to obtain a bacterial concentration of approximately 2×10^6^ CFU mL^-1^. ΦBp-AMP1 was added to these three sub-cultures at an MOI of 1 and the sub-cultures were incubated at 37 °C and 200 rpm. Triplicate aliquots were taken from each infected sub-culture every two hours, serially diluted in LB and plated on LB agar plates. These plates were incubated 37 °C for 24h before counting CFU on each plate to calculate the bacterial concentration at each time point in each sub-culture as the mean and standard error of the mean of biological triplicate each containing technical triplicate. The growth of control uninfected cultures was measured in a similar manner without the addition of phage. Secondly, we measured the change of bacterial density over time during phage infection by measuring the optical density of growing sub-cultures. Briefly, three overnight cultures of *B. thailandensis* were diluted in LB in the wells of a 96 well plate and mixed with phage to obtain a final bacterial concentration of 5×10^5^ CFU mL^-1^ and a final phage MOI of either 10^-4^, 10^-2^, 1 or 10^2^. The plates were the incubated at 37 °C and 200 rpm and the optical density of each well was measured every 10 min via a CLARIOstar plate reader system (BMG) and blank corrected to the optical density measured in wells containing LB medium only. Each condition was tested in 84 technical replicates obtained from biological triplicate.

### Determination of heritable resistance to phage

In order to determine whether heritable resistance to phage had emerged in bacteria from wells where we measured bacterial growth in the presence of phage, we re-inoculated these survivors in wells of a 96-well plate containing LB and phage at a concentration of ∼5×10^5^ PFU mL^-1^. We incubated these plates at 37 °C and 200 rpm for 24 h and measured the optical density as described above. We employed this same approach to determine whether heritable resistance to phage had emerged in bacteria from wells where we measured bacterial growth in the presence of both phage and each of the antibiotics reported in Table 1.

### Determination of colony size

To determine whether phage-resistance comes at a fitness cost in colony size we performed image analysis experiments of bacteria from wells where we measured bacterial growth in the presence of phage. These survivors were plated onto LB agar plates alongside uninfected bacteria from separate control experiments. The plates were then incubated at 37 °C for 72 h and were imaged with a Xiaomi Mi A2 mobile phone camera at 24 h intervals. The colony size was determined via image analysis of 150 colonies for each condition and time point using the ImageJ software.

### Determination of antibiotic minimum inhibitory concentrations and the impact of phage on antibiotic efficacy

The minimum inhibitory concentration (MIC) of each antibiotic employed was determined via the broth dilution method ^79^. Briefly, each antibiotic was dissolved according to the manufacture’s specifications and diluted in a two-fold dilution series in LB medium within wells of a 96-well plate. Stationary *B. thailandensis* E264 bacteria were then added at a final concentration of 5×10^5^ CFU mL^-1^ to each well. The plates were incubated at 37 °C and 200 rpm and after 24 h the optical density of each well was measured via a CLARIOstar plate reader system (BMG) and blank corrected to the optical density measured in wells containing LB medium only. The minimum inhibitory concentration was determined as the minimum concentration of antibiotic for which we measured an optical density value that was less than 10% of the optical density value that we measured in wells containing LB medium and bacteria only. All MIC assays were performed at least in biological and technical triplicate from which mean and standard error of the mean were calculated. Next, to determine the influence of ΦBp-AMP1 on the efficacy of each antibiotic, the experiments above were repeated with the addition of ΦBp-AMP1 at an MOI of 1 (i.e. 5×10^5^ PFU mL^-1^).

### Phage growth in the presence of antibiotics

Overnight cultures of *B. thailandensis* E264 were diluted 1000× in 50 mL LB medium to obtain a bacterial concentration of approximately 2×10^6^ CFU mL^-1^. ΦBp-AMP1 was added to these sub-cultures at an MOI of 1 and the sub-cultures were incubated at 37 °C and 200 rpm. In separate experiments, ΦBp-AMP1 was added at an MOI of 1 together with either ampicillin or ciprofloxacin or trimethoprim at 0.25× their respective MIC. 1 mL triplicate aliquots were taken from the infected sub-cultures at two-hour intervals for twenty-four hours and the phage propagation over time was monitored by determining plaque forming units per millilitre via the double agar overlay technique above. Each experiment was then performed in biological triplicate.

### Determination of time-dependent bacterial growth in the presence of antibiotic and phage

Overnight cultures of *B. thailandensis* E264 were diluted to a concentration of ∼5×10^5^ CFU mL^-1^ in LB medium in wells of a 96-well plate, together with either ΦBp-AMP1 at an MOI of 1, or levofloxacin, cefaclor or trimethoprim at 0.25× their respective MIC, or both ΦBp-AMP1 at an MOI of 1 and either levofloxacin, cefaclor or trimethoprim at 0.25× their respective MIC. Each plate was then incubated for 24 h in a CLARIOstar plate reader (BMG) at 37 °C and 200 rpm, measuring optical density at λ=600nm (OD_600_) every 30 min. Each experiment was repeated in biological and technical triplicate from which we calculated mean and standard error of the mean of each measurement.

### Bacterial killing assays

Bacterial killing assays were performed as previously reported ^39^. Briefly, overnight cultures of *B. thailandensis* E264 were diluted 1000× in 50 mL fresh LB growth medium and infected with either solely ciprofloxacin (at a concentration between 0.625× and 8× the MIC), or a combination of ciprofloxacin and phage at MOI = 1. The cultures were incubated at 37 °C and 200 rpm for 24 h after which the colony count (CFU mL^-1^) was determined as reported above. We quantified bactericidal efficacy of each treatment as the ratio of the colony counts in untreated control experiments over the colony counts measured in each treatment.

### Determination of heritable resistance to antibiotics

In order to determine whether heritable resistance to antibiotics had emerged in bacteria from wells where we measured bacterial growth in the presence of phage and antibiotics, we re-inoculated these survivors in wells of a 96-well plate containing LB and either phage at a concentration of ∼5×10^5^ PFU mL^-1^, or the antibiotic employed at a concentration in range 0.125-128 µg ml^-1^, or both phage at a concentration of ∼5×10^5^ PFU mL^-1^ and the antibiotic employed at a concentration in range 0.125-128 µg ml^-1^. We incubated these plates at 37 °C and 200 rpm for 24 h and measured the optical density as described above.

### Determination of the impact of antibiotics on plaque size

To determine whether phage plaque sizes may be influenced by the presence of antibiotics, we implemented a previously reported protocol ^28^. Briefly, LB top agar was melted in a microwave and allowed to cool below 40 °C. An aliquot of an overnight *B. thailandensis* E264 culture was added to the melted top agar at a volume: volume concentration of 1:50 alongside phage at a concentration of 10^1^ PFU mL^-1^ and either ciprofloxacin, trimethoprim or ampicillin at 0.5× and 0.125× of their respective monotherapy MICs. The resulting mixture was added to the top agar. The top agar was then well mixed by inversion, plated evenly onto LB agar plates (10 g/L, 1.5% Agar) and incubated for 24 h at 37 °C. The plates were then imaged using a Xiaomi Mi A2 mobile phone camera. The plaque size was determined via image analysis of 120 plaques for each condition in biological triplicate using the ImageJ software.

### Determination single-cell morphology and ciprofloxacin accumulation

To measure the size and morphology of individual bacterial during phage-monotherapy or antibiotic-phage combination therapy, we deployed the microfluidic mother machine device as previously described ^80^. Briefly, the device consists of a central channel that measures 25 µm and 100 µm in height and width, respectively, and six thousand lateral side channels, each 1 µm in width and height and 25 µm in length ^81^. Bacteria were prepared by pelleting an overnight culture of *B. thailandensis* for 15 minutes at 3000 *g*. The resulting bacterial pellet was resuspended at a previously optimised nominal OD_600_ of 50 in medium obtained by double filtering the supernatant from the spun down culture using 0.22 µm filters ^82,83^. These bacteria were then introduced in the central channel of the mother machine device from where they reached the lateral channel at an average concentration of one bacterium per channel ^84^. Next, the device was mounted on an inverted microscope (IX73 Olympus, Tokyo, Japan) located in a temperature-controlled chamber kept at 37 °C ^85^. Fluorinated ethylene propylene tubing (1/32” × 0.008”) was connected to the device as inlet and outlet tubes further connected to a computerised pressure-based flow control system (MFCS-4C, Fluigent) ^86^. Next, LB medium only or LB medium containing 2×10^8^ PFU mL^-1^ phage or LB medium containing 2×10^8^ PFU mL^-1^ phage and 0.125× MIC ciprofloxacin was continuously supplied in the device at a constant flow rate of 100 µl/h. Simultaneously, bright field images of 20 areas of the mother machine, each containing 23 lateral channels were acquired at 2 min intervals via a 60× 1.2 N.A. objective (UPLSAPO60XW, Olympus) and an sCMOS camera with an exposure time of 0.01 s (Zyla 4.2, Andor, Belfast, United Kingdom) controlled via Labview ^87^. This microfluidics-microscopy platform was also used to quantify the accumulation of ciprofloxacin as previously reported ^88,89^. Briefly, LB medium containing a fluorescent ciprofloxacin derivative, i.e. ciprofloxacin-nitrobenzoxadiazole (ciprofloxacin-NBD) or ciprofloxacin-NDB and 2×10^8^ PFU mL^-1^ phage was continuously supplied in the device at a constant flow rate of 100 µl/h. In both cases ciprofloxacin-NBD was supplied at a concentration of 32 µg ml^-1^. Bright field images were acquired as described above together with corresponding fluorescence images acquired by exposing the bacteria for 0.03 s to the blue excitation band of a broad-spectrum LED (CoolLED pE300white, power = 8 mW at the sample plane, Andover, UK) via a FITC filter. All images were analysed using the ImageJ software as previously described ^90^.

### Determination of mutations underpinning resistance to phage

In order to determine the mutations underpinning resistance to phage in bacteria from wells where we measured bacterial growth in the presence of phage, we plated these survivors on LB agar plates, incubated these plates at 37 °C for 24h and shipped the plates to MicrobesNG. MicrobesNG performed DNA isolation and ull genome sequencing using 2×250 bp paired end sequencing with a minimum coverage of 30× on the Illumina HiSeq sequencer. Analysis of the sequenced genomes was performed using the *breseq* pipeline ^91^.

### Comparative bacterial and phage transcriptomic analysis

RNA isolation, library preparation, sequencing, and transcriptomic data processing was performed as previously reported ^92^. Briefly, RNA isolation was performed using the RNeasy Mini kit (QIAGEN), according to the manufacturer’s specifications. Bacteria were grown in triplicate for 4h in flasks containing 50 mL LB, or LB containing phage at an MOI of 1, or LB containing ciprofloxacin at 0.125× MIC, or LB containing both phage at an MOI of 1 and ciprofloxacin at 0.125× MIC. DNA removal during extraction was carried out using RNase-Free DNase I (Qiagen). RNA concentration and quality were measured using Qubit 1.0 fluorometer (ThermoFisher Scientific) and 2200 TapeStation (Agilent), respectively, and only samples with an RNA integrity number above 8 were sequenced using Illumina NovaSeq 6000. Transcript abundance was quantified using Salmon for each gene in all samples. Subsequent differential analysis was performed using DEseq2 in R software to quantify the log2 fold change in transcript reads for each gene and compared across the four different experimental conditions. Significantly differentially expressed genes were defined as having a log2 fold change greater than 1 and a p-value adjusted for false discovery rate of < 0.05. Gene Set Enrichment Analysis was performed using the clusterProfiler package for R ^93^. Enrichments in terms belonging to the “Biological process”, “Molecular function” or “Cellular component” ontology were calculated by ranking the genes by differential expression and calculating an Enrichment Score (a weighted Kolmogorov–Smirnov-like statistic) for each ontology term. P values were adjusted for false discovery by using the method of Benjamini and Hochberg ^94^. Finally, the lists of significantly enriched terms were simplified to remove redundant terms, as assessed via their semantic similarity to other enriched terms, using clusterProfiler’s simplify function.

### Statistical non-linear regression model

We developed a statistical non-linear regression model to fit the experimental data in order to estimate treatment output and the associated uncertainty also for treatment conditions that we did not investigate experimentally. Since the distribution of bacterial growth values (in terms of OD_600_) in the presence of phage was skewed, we fitted our data with a Gamma distribution function. Moreover, to account for inaccuracies in optical density measurements via a plate reader, we assumed any measurement below an optical density of 0.05 to be equal to ‘zero’. Hence, the data had a non-trivial number of zero values, which could not be captured by a conventional gamma distribution. To accommodate this, we included a parameter which controls the probability of an experiment returning an optical density equal to zero. This two-part distribution strategy resulted in what is often called a “hurdle model” ^95^. This distribution then had three parameters, one that represented the probability of measuring a value of zero, and two representing the gamma distribution, in our case parameterised to have a mean parameter and a shape parameter. We assumed that these three parameters could change with the antibiotic concentration and with the phage concentration. We also assumed these relationships to be potentially non-linear. In other words, we assumed the probability of observing a value of zero, the average non-zero observation, and the spread of non-zero observations all depended on the antibiotic and phage concentrations in a complex manner. As such, we modelled these parameters using cubic regression tensor product smoothing splines ^96^. Additionally, the spline knot locations for the two gamma distribution parameters were constrained to only the points where a non-zero observation was made. To fit this model, and estimate the various parameters, including splines, we used Markov Chain Monte Carlo (MCMC), using the brms R package ^97^, with default options used for the priors. This provided us with posterior distributions for all the unknown parameters, allowing us to make probabilistic statements and provide full uncertainty estimates. To better capture the uneven measurement intervals of empirical data and because we were interested in the relationships between antibiotic and phage concentrations, it was convenient to apply a log2 transform to the antibiotic concentration and a log10 transformation to the phage concentration values. To avoid numerical issues when the antibiotic or phage concentrations were equal to 0, we added a small value to each zero before applying the transformation (0.01 for the antibiotic data, and 1e^-10^ for the phage measurement data). The model allowed us to continuously predict bacterial growth in terms of optical density and the associated uncertainty for both treatment concentrations that were experimentally investigated as well as treatment concentrations that were not experimentally investigated.

## Data availability

All data generated or analysed during this study are included in this published article and its supplementary information files.

## Acknowledgments

This work was supported by the BBSRC through a grant awarded to S.P., K.T.A. and U.L. (BB/V008021/1). S.K. was supported by a QUEX PhD studentship awarded to S.P., M.A.T.B. and K.T.A. KTA and E.B. gratefully acknowledge the financial support of the EPSRC (EP/T017856/1). K.C., K.B. and S.H. were supported by the Defence Science and Technology Laboratory. This project utilised equipment funded by a Wellcome Trust Institutional Strategic Support Fund (WT097835MF), a Wellcome Trust Multi-User Equipment Award (WT101650MA) and a BBSRC LoLa award (BB/K003240/1). The funders had no role in study design, data collection and analysis, decision to publish, or preparation of the manuscript.

## Author contributions

Conceptualization, S.P.; methodology, S.K., U.L., E.B. and E.L.A.; formal analysis, S.K., U. L., E.B. and S.P; generation of figures, S.K., U.L., E.B. and S.P; investigation, S.K., U.L., K.C., E.B., E.L.A., P.O.N., A.F., A.A. E.G., S.K., K.B., S.V.H., K.T.A., M.A.T.B. and S.P.; resources, K.T.A., M.A.T.B. and S.P.; data curation, S.K. and S.P., writing – original draft, S.K. and S.P.; writing – review & editing, S.K., U.L., K.C., E.B., E.L.A., P.O.N., A.F., A.A., E.G., S.K., K.B., S.V.H., K.T.A., M.A.T.B. and S.P.; visualization, S.K., U.L. and S.P.; supervision, S.P., M.A.T.B. and K.T.A.; project administration, S.P.; funding acquisition, S.P., M.A.T.B. and K.T.A..

