## Supporting information for "Phage-induced efflux down-regulation boosts antibiotic efficacy"

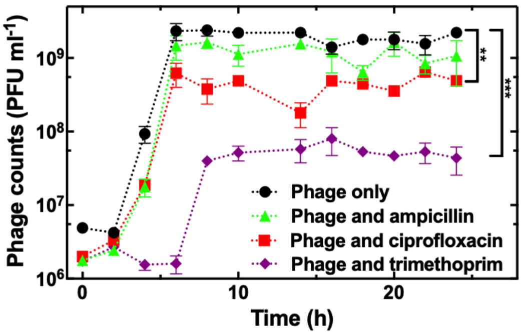


**Figure S1 Phage amplification in the presence of sub-inhibitory concentrations of antibiotics**. Temporal dependence of phage counts from stationary phase *B. thailandensis* cultures incubated in LB medium only (black circles), or in LB medium containing ampicillin (green triangles), ciprofloxacin (red squares) or trimethoprim (purple diamonds) at 0.25× their respective MIC values. In all cases the initial bacterial inoculum was $2\times10^6$ CFU ml$^{-1}$ and the starting concentration of phage was $2\times10^6$ PFU ml$^{-1}$. Symbols and error bars are means and standard errors of the mean of phage count measurements from biological and technical triplicates. Very small error bars cannot be visualised due to overlap with the datapoints. Dashed lines are guides-for-the-eye. **** indicate a p-value < 0.0001.

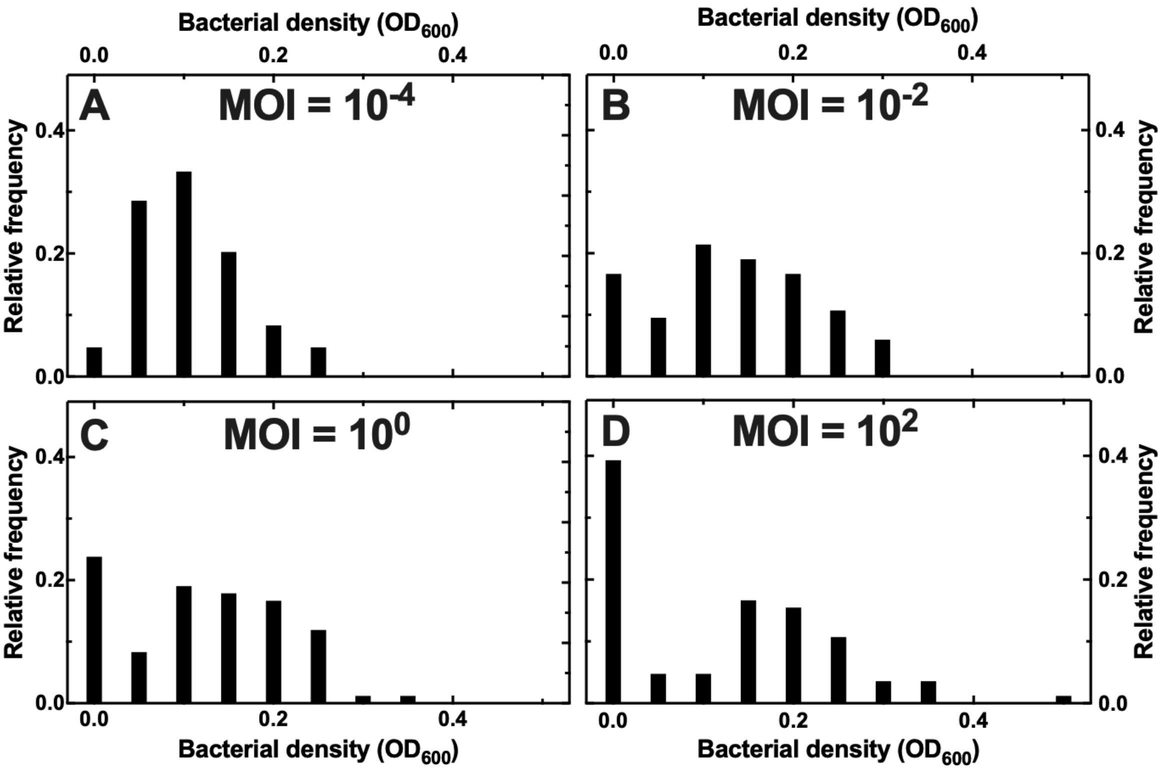


**Figure S2 Dependence of bacterial density on phage MOI**. Distribution of *B. thailandensis* density values, measured in $OD_{600}$, after 24 h exposure to phage at an MOI of (A) $10^{-4}$, (B) $10^{-2}$, (C) $10^{0}$ or (D) $10^{2}$. For each condition, bacterial density measurements were carried out in 84 independent micro-cultures from biological triplicates.

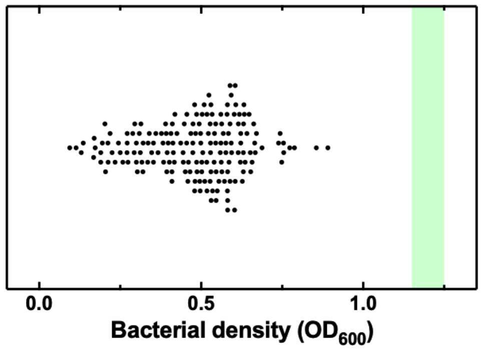


**Figure S3 *B. thailandensis* growth in the presence of phage at 25 °C**. Bacterial density measurements after 72 h incubation at 25 °C in the presence of phage at an MOI of 1. Each black circle represents a bacterial density value performed on one of 84 technical micro-culture replicates from biological triplicates. The green vertical band represents the mean and standard error of the mean of corresponding bacterial density measurements in the absence of phage.

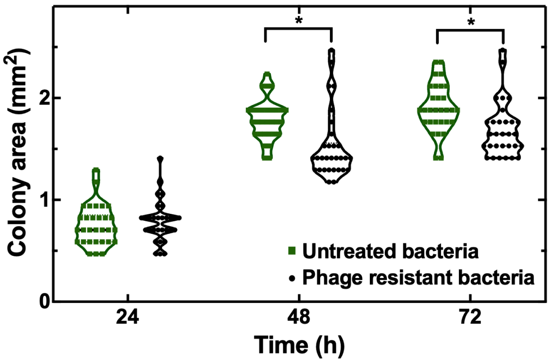


**Figure S4 Colony growth of phage resistant *B. thailandensis*.** Temporal dependence of the area of colonies of untreated *B. thailandensis* (green squares) and of phage resistant mutants (black circles). Each symbol represents the measurement of the area of one of 30 independent colonies from biological triplicates * indicates a p-value < 0.05.

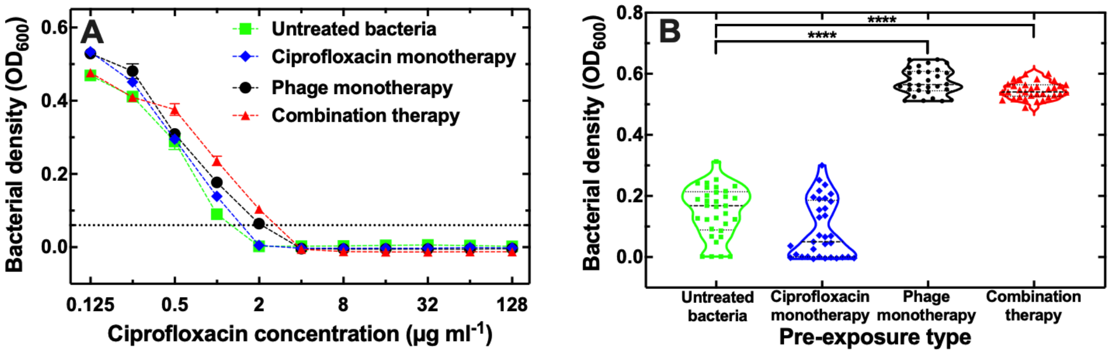


**Figure S5 Resistance to phage and ciprofloxacin.** (A) Dependence of bacterial density, measured in $OD_{600}$, on the concentration of ciprofloxacin for *B. thailandensis* that had not undergone any previous therapy (green squares), *B. thailandensis* that had undergone a 24 h monotherapy with ciprofloxacin at 0.125× its MIC (blue diamonds), *B. thailandensis* that had undergone a 24 h monotherapy with phage at an initial MOI of 1 (black circles), or *B. thailandensis* that had undergone a 24 h combination therapy with ciprofloxacin at 0.125× its MIC and phage at an initial MOI of 1 (red triangles). Symbols and error bars are means and standard errors of the means of bacterial density measurements obtained from biological triplicates each containing five technical micro-culture replicates. Very small error bars cannot be visualised due to overlap with the datapoints. Dashed lines are guides-for-the-eye. The horizontal dashed line represents 10% of the bacterial density value measured after 24 h incubation in LB medium in the absence of ciprofloxacin and phage. (B) Bacterial density, measured in $OD_{600}$, after 24 h exposure to phage at an initial MOI of 1 for *B. thailandensis* that had not undergone any previous therapy (green squares), *B. thailandensis* that had undergone a 24 h monotherapy with ciprofloxacin at 0.125× its MIC (blue diamonds), *B. thailandensis* that had undergone a 24 h monotherapy with phage at an initial MOI of 1 (black circles), or *B. thailandensis* that had undergone a 24 h combination therapy with ciprofloxacin at 0.125× its MIC and phage at an initial MOI of 1 (red triangles). Each symbol represents a bacterial density measurement obtained from one of 30 technical micro-culture replicates from biological triplicates. The median and quartile of each distribution are indicated as black dashed and dotted lines, respectively.

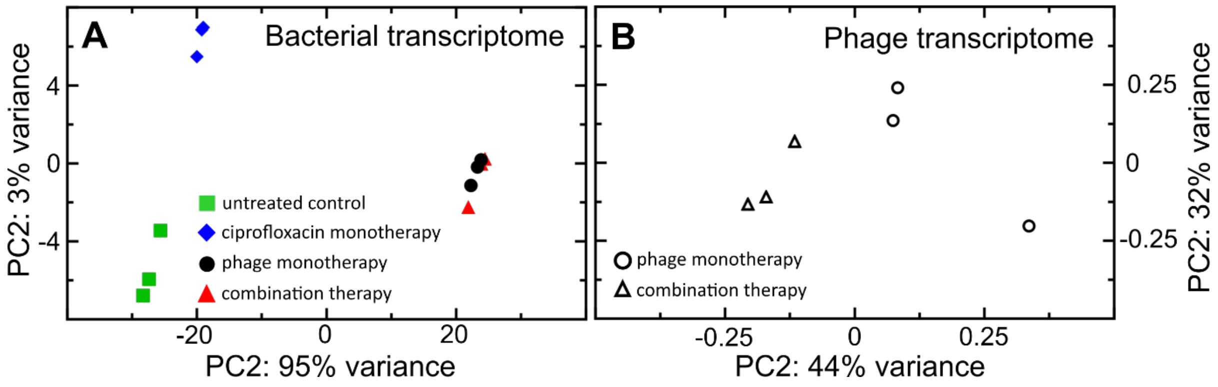


**Figure S6 Principal component analysis of bacterial and phage transcriptomes.** (A) Principal component analysis of replicate transcriptomes of stationary phase *B. thailandensis* incubated for 4 h either in LB medium only (green squares), or in LB medium containing ciprofloxacin at 0.125× its MIC (blue diamonds), or LB medium containing phage at an MOI of 1 (black circles), or in LB medium containing both ciprofloxacin at 0.125× its MIC and phage at an MOI of 1 (red triangles). (B) Corresponding principal component analysis of replicate transcriptomes of phage ΦBp-AMP1 after 4 h incubation with stationary phase *B. thailandensis* in LB medium only (open circles) or in LB medium containing ciprofloxacin at 0.125× its MIC (open triangles). Note the narrower x-axis range with respect to Figure S6A because of the largely overlapping phage transcriptomes.

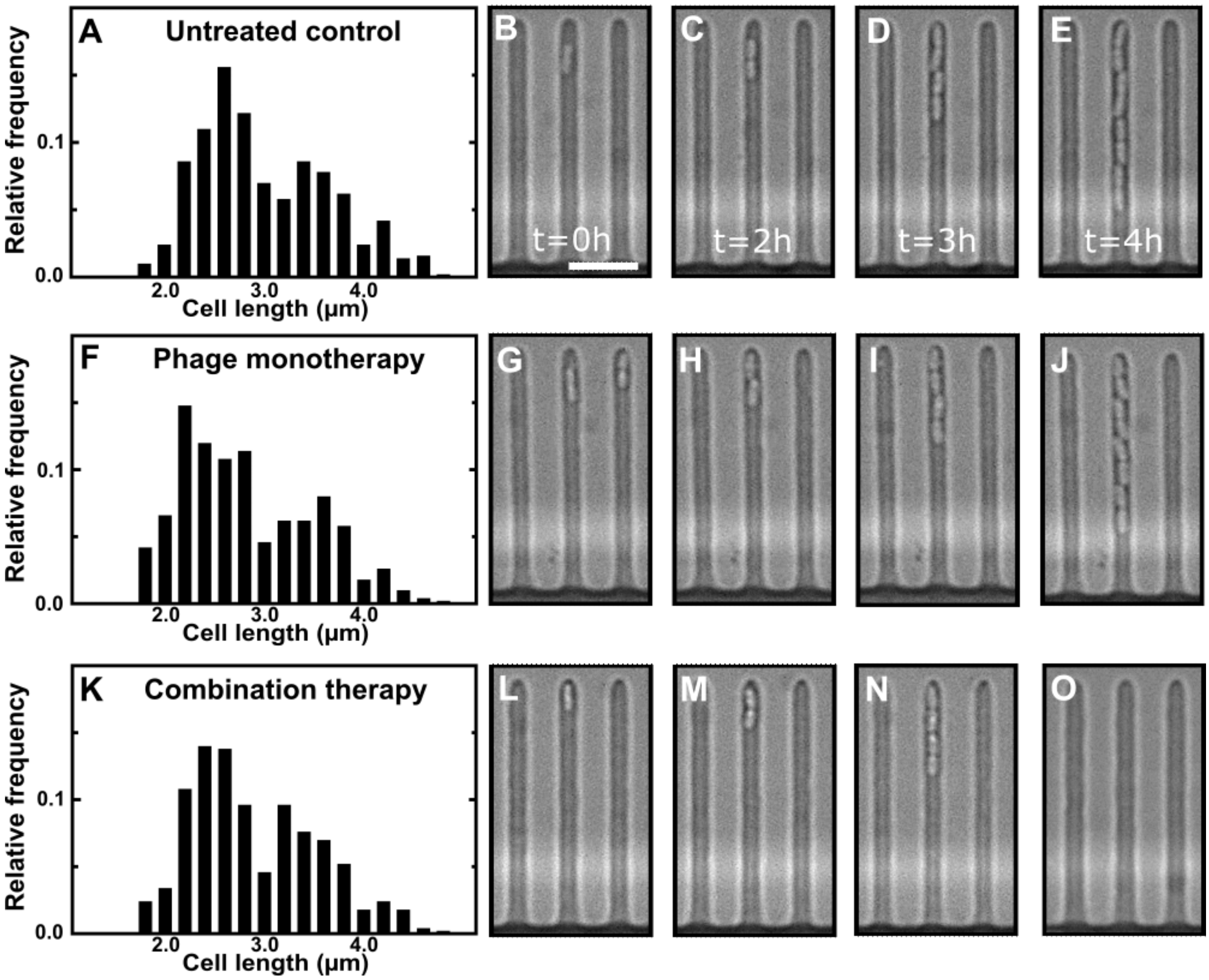


**Figure S7 Single-cell morphology.** Distribution of single-cell lengths for (A) *B. thailandensis* incubated in LB medium only, (F) *B. thailandensis* incubated in LB medium containing phage at a concentration of $2\times10^8$ PFU $ml^{-1}$ and (K) *B. thailandensis* incubated in LB medium containing both phage at a concentration of $2\times10^8$ PFU $ml^{-1}$ and ciprofloxacin at 0.125× its MIC. Each distribution contains 500 single-cell length measurements carried out on bacteria hosted in different microfluidic compartments from different biological triplicate experiments over a period of 9 h exposure to each condition. Corresponding representative microscopy images are reported in (B-E), (G-J) and (L-O), respectively. Scale bar: 5 µm.

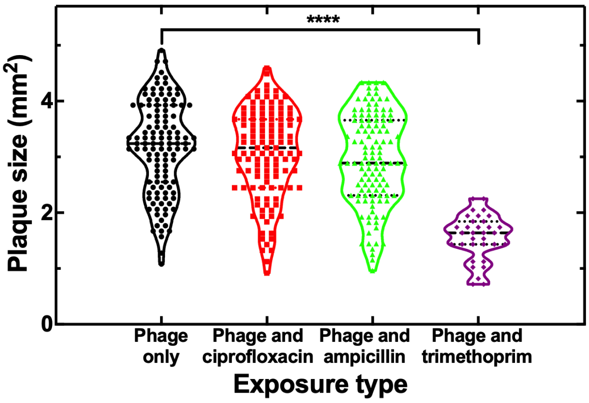


**Figure S8 Plaque size in the presence of sub-inhibitory concentrations of antibiotics.** Distribution of sizes of plaques formed ΦBp-AMP1 when plated on *B. thailandensis* alone (black circles), or in the presence of ciprofloxacin (red squares), ampicillin (green triangles) or trimethoprim (purple diamonds) at 0.125× their respective MIC values. Each symbol reports a plaque size value measured on one of 150 plaques for each condition. Dashed and dotted horizontal lines represent the median and quartiles, respectively, of each distribution. **** indicates a p-value < 0.0001. Similar data were obtained when these antibiotics were used at 0.5× their respective MIC values.

| Gene | Product | Annotation | Mutation | Resistance |
|---|---|---|---|---|
| *BTH_RS14320* → | O-acetyl-ADP-ribose | V38G (GTC→GGC) | T→G | Low |
| *wbiB* → | dTDP-L-rhamnose | coding (415/1092 nt) | $(CCGAGCAG)_{1\rightarrow 2}$ | Low |
| *wbiB* → | dTDP-L-rhamnose | coding (473/1092 nt) | +G | Low |
| *BTH_RS19740* → | O-antigen methyl | S476P (TCC→CCC) | T→C | High |
| *BTH_RS19745* → | Glycosyltransferase | I231S (ATC→AGC) | T→G | High |
| *BTH_RS19745* → | Glycosyltransferase | P521L (CCG→CTG) | C→T | High |
| *BTH_RS19765* → | Polysaccharide | H478Y (CAT→TAT) | C→T | High |
| *BTH_RS21230* →/- | Hypothetical Protein - | intergenic (+131/-129) | $(AAGGGCTC)_{7\rightarrow 8}$ | Low |
| *mlaE* ← | ABC transporter | *256Y (TAA→TAC) | T→G | Low |
| *rfbA* → | Thymidylyltransferase | coding (675/894 nt) | Δ1 bp | Low |

**Table S1 Unique mutations identified in representative low- and high-resistance mutants.** Gene, gene product, annotation, unique mutation and level of resistance to phage measured in five representative low- and high-resistant mutants.

| Level of resistance | *Burkholderia phage* | | | | | *Ralstonia phage* |
|---|---|---|---|---|---|---|
| | **ΦBp-AMP1** | **ΦE12-2** | **ΦE125** | **KL3** | **vB_BmuP_KL4** | **RsoM1USA** |
| | NC_047743 | NC_009236 | NC_003309 | NC_015266 | NC_047958 | NC_049432 |
| **High** | 0 | 150 | 150 | 20 | 0 | 0 |
| | 0 | 150 | 150 | 0 | 0 | 20 |
| | 0 | 150 | 150 | 0 | 0 | 20 |
| | 0 | 150 | 150 | 40 | 0 | 20 |
| | 0 | 150 | 150 | 40 | 0 | 20 |
| **Low** | 0 | 150 | 140 | 40 | 0 | 20 |
| | 0 | 150 | 150 | 0 | 0 | 20 |
| | 0 | 150 | 150 | 0 | 0 | 20 |
| | 0 | 150 | 150 | 0 | 40 | 20 |
| | 0 | 150 | 150 | 0 | 0 | 20 |
| **Untreated Control** | 0 | 150 | 150 | 0 | 0 | 20 |
| | 0 | 150 | 150 | 0 | 40 | 20 |
| | 0 | 150 | 150 | 40 | 0 | 20 |
| | 0 | 150 | 150 | 40 | 0 | 20 |
| | 0 | 150 | 150 | 40 | 0 | 20 |

**Table S2 Phage within uninfected *B. thailandensis* cultures**. Score of integrity for *Bulkholderia* phage ΦBp-AMP1, ΦE12-2, ΦE125, KL3 and vB_BmuP_KL4, and the *Ralstonia* phage RsoM1USA measured from 5 representative high-resistant mutant cultures, 5 representative low-resistant mutant cultures and 5 untreated control cultures, all in the absence of externally added phage. Detected phage sequences were given a score of integrity via PHASTER, with a maximum value of 150, scores above 90 were considered as intact prophages (blue shaded tabs), scores below 70 were considered as incomplete (red shaded tabs).

| Class | Molecule | Predicted additive range ($\mu g\ ml^{-1}$) |
|---|---|---|
| Quinolones | Nalidixic Acid | 2 - 16 |
| | Ciprofloxacin | 0.06 - 2 |
| | Ofloxacin | 0.5 - 4 |
| | Levofloxacin | 0.5 - 4 |
| | Finafloxacin | 0.06 - 2 |
| | Moxifloxacin | 0.06 - 2 |
| β-Lactams | Amoxicillin | 12 - 32 |
| | Ampicillin | 12 - 32 |
| | Cefaclor | 64 - 256 |
| | Ceftazidime | 0.5 - 2.5 |
| | Meropenem | 1.25 - 4 |
| Tetracycline | Doxycycline | 1 - 4 |
| | Tetracycline | 2 - 8 |

**Table S3 Additivism between phage and antibiotics.** Antibiotic class, antibiotic molecule and range of antibiotic concentrations for which our model predicted additivism between phage and each antibiotic based on our experimental MIC values recorded for monotherapy and combination therapy using each antibiotic. Interactions between phage and antibiotics were considered to be additive if the probability that combination therapy was more effective in inhibiting bacterial growth than phage and antibiotic monotherapies was above 95%.

**Supplementary files:**

**Supplementary file 1. Differential expression of *B. thailandensis* genes after ciprofloxacin monotherapy relative to untreated control *B. thailandensis*.** Transcript base mean, log2 fold change, standard error of log2 fold change, p-value, adjusted p-value, gene identifier, gene and gene product for each of 4920 differentially expressed *B. thailandensis* genes, identified via Illumina sequencing, between ciprofloxacin monotherapy relative to untreated control *B. thailandensis*.

**Supplementary file 2. Differential expression of *B. thailandensis* genes after phage monotherapy relative to untreated control *B. thailandensis*.** Transcript base mean, log2 fold change, standard error of log2 fold change, p-value, adjusted p-value, gene identifier, gene and gene product for each of 5229 differentially expressed *B. thailandensis* genes, identified via Illumina sequencing, between phage monotherapy relative to untreated control *B. thailandensis*.

**Supplementary file 3. Differential expression of *B. thailandensis* genes after combination therapy relative to untreated control *B. thailandensis*.** Transcript base mean, log2 fold change, standard error of log2 fold change, p-value, adjusted p-value, gene identifier, gene and gene product for each of 5215 differentially expressed *B. thailandensis* genes, identified via Illumina sequencing, between combination therapy relative to untreated control *B. thailandensis*.

**Supplementary file 4 Gene ontology enrichment analysis of differentially expressed *B. thailandensis* genes between ciprofloxacin monotherapy and untreated control bacteria.** Identifier, description, number of genes contained, enrichment score, normalized enrichment score, p-value, adjusted p-value, q-value, rank, leading edge and identifier of each gene contained for each significantly enriched biological process, molecular function or cellular component in the comparison between ciprofloxacin monotherapy and untreated control bacteria.

**Supplementary file 5 Gene ontology enrichment analysis of differentially expressed *B. thailandensis* genes between phage monotherapy and untreated control bacteria.** Identifier, description, number of genes contained, enrichment score, normalized enrichment score, p-value, adjusted p-value, q-value, rank, leading edge and identifier of each gene contained for each significantly enriched biological process, molecular function or cellular component in the comparison between phage monotherapy and untreated control bacteria.

**Supplementary file 6 Gene ontology enrichment analysis of differentially expressed *B. thailandensis* genes between combination therapy and untreated control bacteria.** Identifier, description, number of genes contained, enrichment score, normalized enrichment score, p-value, adjusted p-value, q-value, rank, leading edge and identifier of each gene contained for each significantly enriched biological process, molecular function or cellular component in the comparison between combination therapy and untreated control bacteria.

**Supplementary file 7. Differential expression of ΦBp-AMP1 genes after combination therapy relative to phage monotherapy.** Transcript base mean, log2 fold change, standard error of log2 fold change, p-value, adjusted p-value, gene identifier, gene and gene product for each of 41 differentially expressed ΦBp-AMP1 genes, identified via Illumina sequencing, between combination therapy relative to phage monotherapy.
